# An integrative approach leads to the discovery of a novel anti-leukemic peptide from human milk

**DOI:** 10.1101/2021.03.07.434306

**Authors:** Wararat Chiangjong, Jirawan Panachan, Thitinee Vanichapol, Nutkridta Pongsakul, Pongpak Pongphitcha, Teerapong Siriboonpiputtana, Tassanee Lerksuthirat, Pracha Nuntnarumit, Sarayut Supapannachart, Chantragan Srisomsap, Jisnuson Svasti, Suradej Hongeng, Somchai Chutipongtanate

## Abstract

Chemotherapy in childhood leukemia is associated with late morbidity in leukemic survivors, while certain patient subsets are relatively resistant to standard chemotherapy. It is therefore important to identify new agents with sensitivity and selectivity towards leukemic cells, while having less systemic toxicity. Peptide-based therapeutics has gained much attention during the last few years. Here, we used an integrative workflow combining mass spectrometric peptide library construction, *in silico* anticancer peptide screening, and *in vitro* leukemic cell studies to discover a novel anti-leukemic peptide having 3+charges and alpha-helical structure, namely HMP-S7, from human breast milk. HMP-S7 showed cytotoxic activity against four distinct leukemic cell lines in a dose-dependent manner but had no effect on solid malignancies or representative normal cells. HMP-S7 induced leukemic cell death by penetrating the plasma membrane to enter the cytoplasm and cause leakage of lactate dehydrogenase, thus acting in a membranolytic manner. Importantly, HMP-S7 exhibited anti-leukemic effect against patient-derived leukemic cells *ex vivo*. In conclusion, HMP-S7 is a selective anti-leukemic peptide with promise which requires further validation in preclinical and clinical studies.

**Teaser:** *In silico* screening of naturally occurring human milk peptides discovers a new anticancer peptide that kills leukemic cells *in vitro* and *ex vivo*.

## MAIN TEXT

### Introduction

Cancer is a significant cause of death in children and adolescents during the last few years in Asia, Central and South America, Northwest Africa, and the Middle East (*1*). Hematologic malignancies, particularly acute lymphocytic leukemia (ALL), are predominant, accounting for 30% of childhood cancers (*2*). Optimization of chemotherapeutic regimens during the past decades has resulted in more than 90% remission rate in childhood ALL (*3*). However, the burden of late morbidity due to chemotherapeutic treatments, which occur in two-thirds of pediatric cases (*4*), have become important considerations as the number of long-term leukemic survivors increases (*5*). Moreover, specific subsets of pediatric ALL are relatively resistant to standard chemotherapy and have high risk of relapse (*6*). These challenges stress the need to further improve current treatment, while identifying new agents with sensitivity and selectivity toward leukemic cells with little to no toxicity to normal cells.

Peptide-based drugs or anticancer peptides can provide a new strategy for cancer treatment. Although the exact mechanisms and selectivity criteria have yet to be elucidated, anticancer peptides may have oncolytic effects depending on peptide characteristics and target membrane properties in determining selectivity and toxicity (*7*). Cancer cells have highly negative transmembrane potential from anionic molecules on the surface, greater membrane fluidity, and more abundant microvilli (increasing outer surface area). In contrast, normal cells are electrically neutral (*8–11*). The negative charges on the cancer cell membrane attract positively charged peptides which will disturb membrane stability, causing the loss of electrolytes and cell death (*8*), while the high cholesterol content of the normal cell membrane can protect cell fluidity and block cationic peptide entry (*12*). Common strategies for anticancer peptide discovery include: a) activity-guided purification from biological/natural products (*13*); b) examination of antimicrobial peptides for cancer sensitivity (*14, 15*). The former approach tends to deliver positive results but is associated with labor-intensive and time-consuming processes. The latter strategy is relatively cost- and time-effective but still tends to be limited to antimicrobial peptides known *a priori*. Investigation of anticancer peptides has progressed slowly in the past decades, indicating gaps for improvement, particularly on the selection of peptide sources, screening methods, and downstream analyses. Human milk is a promising source of therapeutic peptides. Bioactive milk peptides are released from source proteins by enzymatic hydrolysis, fermentation with a proteolytic starter culture, and proteolysis with proteolytic microorganisms (*16, 17*). Although human milk has rarely been investigated for anticancer activities, studies have showed human milk-derived peptides could exhibit various biological effects such as antimicrobial (*18, 19*) and immunomodulatory activities (*20*). Human milk-derived beta-casein fragments have been studied for immunomodulation, antibacterial, antioxidant, opioid agonist, antihypertensive activities, as well as cell proliferation of human preadipocytes (*21*). These bioactive peptides have different amino acid compositions and sequences, and the peptide length can vary from two to twenty amino acid residues (*22*). It was anticipated that human milk also contains anticancer peptides, as found with bovine milk. PGPIPN hexapeptide of bovine β-casein can inhibit invasion and migration of human ovarian cancer cells (*23*). ACFP, an anti-cancer fusion peptide derived from bovine β-casein and lactoferrin, can inhibit viability and promote apoptosis in primary ovarian cancer cells (*24*). Interestingly, breastfeeding for six months or longer was associated with a 19% lower risk of all childhood leukemia than shorter breastfeeding or none (*25, 26*). Identifying novel anti-leukemic peptides from human milk is important for the future development of non-allergic and non-toxic peptide-based drugs to improve childhood leukemia therapy.

This study aimed to discover a novel anti-leukemic peptide from human milk. We applied a robust workflow integrating the strength of liquid chromatography-tandem mass spectrometry to generate a library of naturally occurring human milk peptides, *in silico* screening based on physicochemical, structural, and predictive anticancer properties, using machine learning algorithms. Potential candidates were selected for functional studies of synthetic peptides to determine anti-leukemic activity and identify the mode of action. By this strategy, a novel anti-leukemic peptide was identified from human milk as the main outcome, and its anti-leukemic activity was successfully validated against four distinct leukemic cell lines *in vitro*, as well as three patient-derived leukemic cells *ex vivo*.

## Results

To separate human milk peptides and test their cytotoxicity to leukemic cells, ten healthy mothers aged between 25 and 36 years old (32.3±3.27 years old) donated breast milk at once for 6-129 days (63.8±38.97 days) after delivery (details in **Table S1)**. After multiple steps of centrifugation to remove cells, lipids, and extracellular vesicles, ultrafiltration (3-kDa cutoff) was used to remove proteins, the small molecules (<3-kDa) were subjected to C18 SPE. The C18 bound peptides were eluted by various acetonitrile concentrations and tested for their cytotoxicity. The conceptual framework of this study is illustrated in **Fig. 1**.

**Fig. 1.**
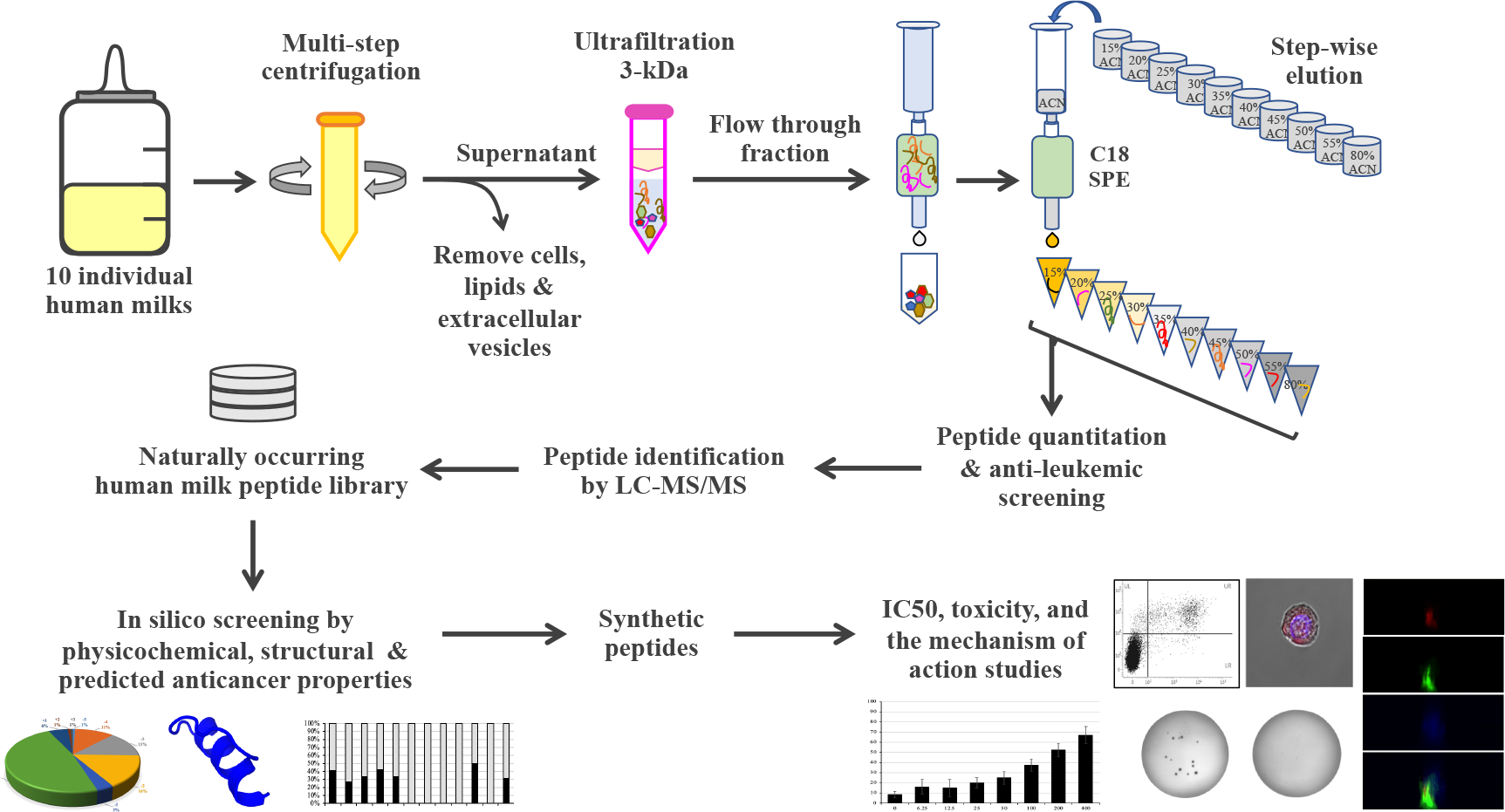
A conceptual framework of the integrative strategy for discovering a novel human milk-derived anti-leukemic peptide. This strategy combines the strengths of mass spectrometry for high-throughput peptide identification, *in silico* screening for prioritizing peptide candidates, and experimental validation for anti-leukemic activities. Abbreviations: LC-MS/MS, liquid chromatography-tandem mass spectrometry; IC_50_, half-maximal inhibitory concentration; SPE, solid-phase extraction.

### Most human milk peptide fractions had cytotoxic effects on leukemic and normal cell lines

After C18 bound peptide elution, the crude milk peptide fraction was tested to observe cytotoxic effects against Jurkat (T lymphoblastic leukemia) and FHs74Int cells (the representative normal intestinal epithelium), respectively (**Fig. 2A**). The crude milk peptides showed a potent cytotoxic effect on Jurkat but had less impact on FHs74Int cells. The coarse milk peptide fraction was then fractionated by C18 SPE with the stepwise acetonitrile (ACN) elution, with the chromatogram showing amounts of peptides at each % ACN (**Fig. 2B**). Peptides eluted with 15%-80% ACN were used to treat Jurkat leukemic cells and FHs74Int normal cells, and cell survival observed as shown in **Fig. 2C**. The relatively low peptide content in the 15%-30% ACN eluted fractions could decrease Jurkat cell survival more than the fraction with the highest peptide content (45% ACN eluted peptide fraction). This indicated that the elimination of Jurkat leukemic cells depended on the specific peptide sequence with hydrophilic properties being more than peptide amount.

**Fig. 2.**
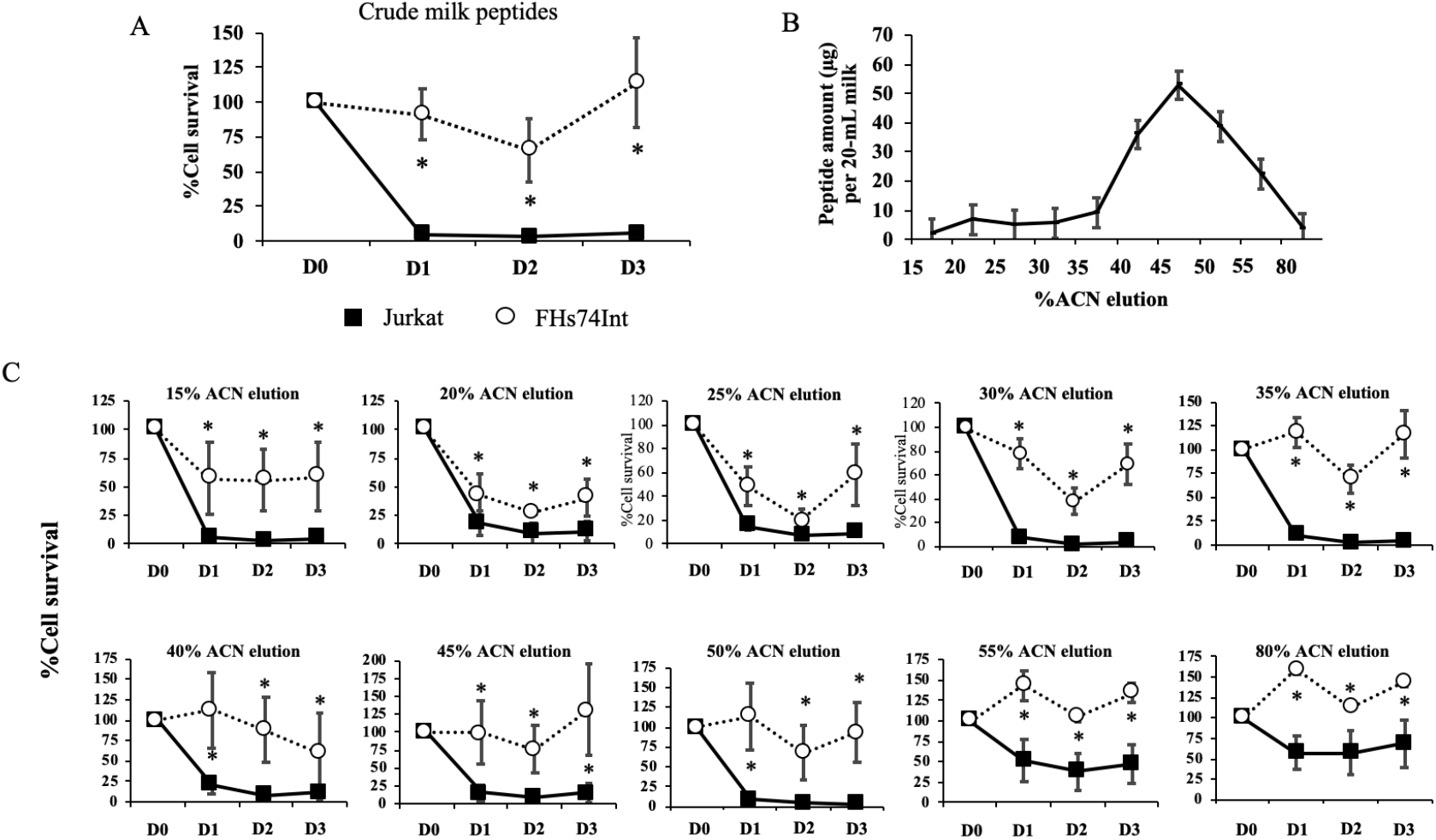
Fractionation of peptides from human milk and cytotoxicity of fractions towards leukemic and normal cells. Ten samples of human milk were divided into 3 pools. Twenty milliliters of each pool were centrifuged at 4°C to remove cells, lipid, and extracellular vesicles. The crude milk peptides obtained from each pool were separately eluted through a <3 kDa cut-off column, and the eluate was loaded to a C18 SPE column. Milk peptides bound to the C18 SPE column were eluted with various concentrations of ACN from 15% ACN to 80% ACN (1 mL each). Eluted fractions of milk peptides were dried using a SpeedVac concentrator, resuspended in culture medium, and then used to treat Jurkat (black square) and FHs74Int cells (white circle), using 3 biological replicates. WST-1 assay was applied to measure cell viability. **A.** % cell survival (mean±SEM) after treatment of cells with crude milk peptides for 1 (D1), 2 (D2) or 3 (D3) days, compared to untreated control (D0); **(B)** Amounts of peptides eluted from C18 SPE column eluted in a stepwise manner using 1 ml each of increasing acetonitrile concentrations of 15%, 20%, 25%, 30%, 35%, 40%, 45%, 50%, 55%, and 80% ACN. Peptides were quantitated by the Bradford method and shown as, mean ± SEM. **(C)** % cell survival (mean±SEM) after treatment of cells with eluates obtained at different %ACN, for 1 day (D1), 2 days (D2) or 3 (D3) days, compared to untreated controls (D0). *; *p*<0.05 comparing to the untreated condition.

### Peptide identification, library construction, and in silico anti-cancer peptide screening

Eleven fractions (from one crude and ten stepwise ACN eluates) of human milk peptides were identified by LC-MS/MS. A total of 142 naturally occurring human milk peptides with unique sequences were collected into the peptide library, ready for further analyses (**Table S2**). The distributions of all identified peptides by fractions, peptide length, p*I*, and the net charge are shown in **Fig. 3A** (upper panel). Overall, most identified natural occurring-milk peptides had fewer than 20 amino acids in length, contained zero or anionic net charge inclined towards acid properties. The most frequently detected pattern was found to be natural peptides with proline-rich sequences (**Fig. S1**). Proline-rich peptides may play important roles in the biological effects of human milk. For example, a ligand containing a proline-rich sequence or a single proline residue may be involved in protein-protein interaction (*27*). Antimicrobial peptides with proline-rich sequences can kill microorganisms by interacting with 70S ribosome and disrupting protein synthesis (*28*).

**Fig. 3.**
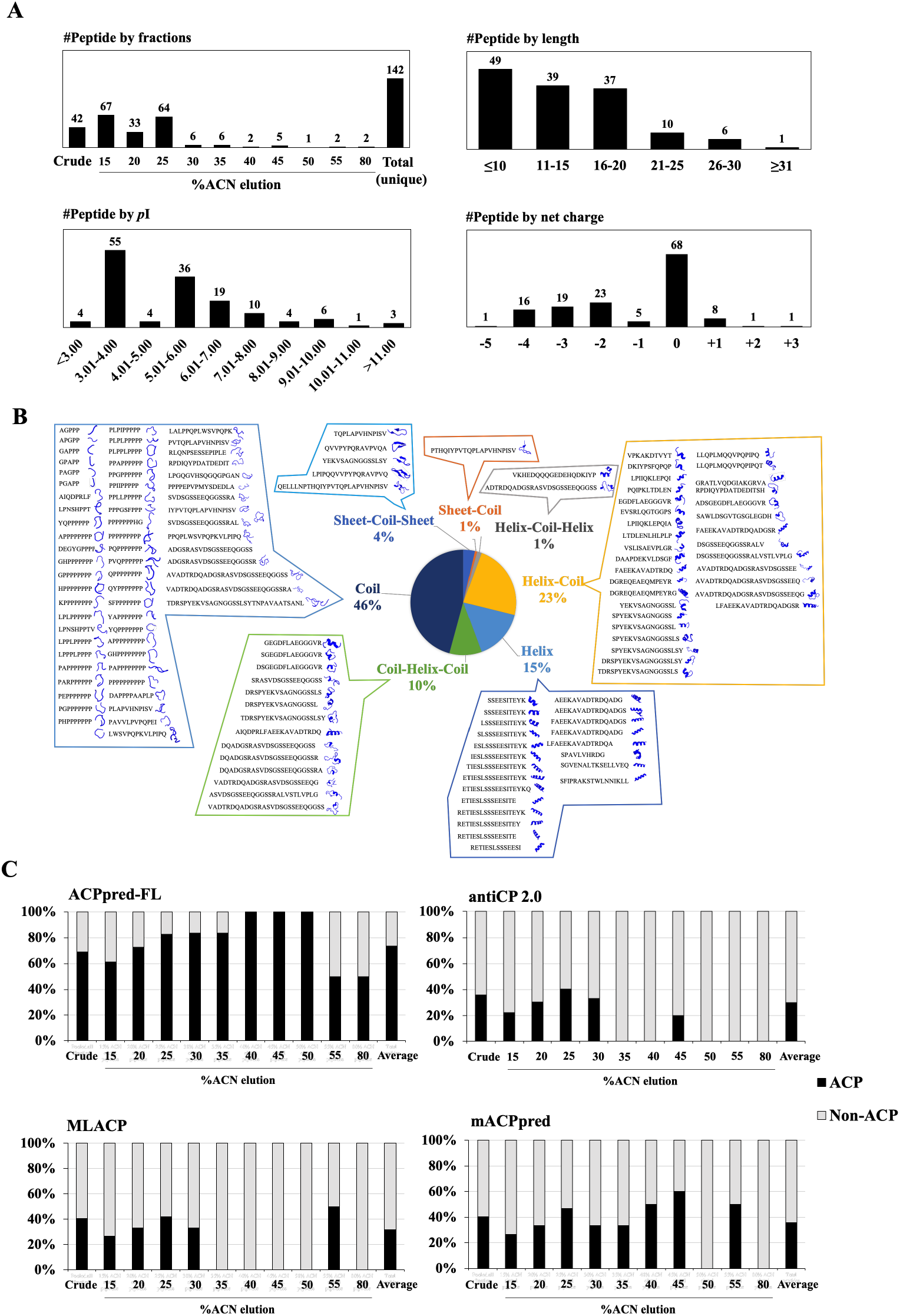
Predicted physicochemical, structural, and machine learning-based anticancer peptide screening of naturally occurring human milk peptides. (**A**) Distribution of the unique human milk peptides identified by LC-MS/MS. (**B**) The distribution and secondary structure of all identified peptides predicted by PEP-FOLD3 software. (**C**) Proportions of predicted ACP vs. non-ACP peptides using four ACP machine learning programs, including ACPpred-FL, antiCP 2.0, MLACP, and mACPpred. The percentage of ACP (black) and non-ACP (gray) peptides was calculated by (number of ACP or non-ACP predictions/number of total identified peptides in each individual fraction) ×100%. Full results of *in silico* ACP screening are provided in **Table S2**.

*In silico* screening of the naturally occurring human milk peptide library was performed using a robust workflow. Firstly, we screened the library by physicochemical properties, i.e., length, *p*I, and net charge (as shown in **Fig. 3A**) since most of the known anticancer peptides are small cationic peptides (commonly 5-30 amino acid length with net charge +2 to +9) that bind to the negative charge of phosphatidylserine and sialic acid on the cancer cell membrane (*12, 29, 30*). Secondly, the secondary structures of the peptides were predicted by the PEP-FOLD3 webserver (**Fig. 3B**). Peptides can form alpha-helix, beta-sheet, and random coils (*31*). However, most of the known anticancer peptides shared the alpha-helix structure (*32*). Thirdly, the identified peptides were predicted for anticancer properties (ACP) using machine learning prediction software. Since ACP prediction software could provide different results depending on the algorithms and training datasets, four online-accessible software were applied in this study, including ACPpred-FL (*33*), antiCP 2.0 (*34*), MLACP (*35*), and mACPpred (*36*) (**Fig. 3C**). The results showed eight human milk peptides and one bovine milk peptide (as a positive ACP control) from various prioritizations based on net charge, secondary structure, and the predicted anticancer property were selected for further analysis (**Table S3)**.

### HMP-S7 killed leukemic cells but not normal cells or solid tumor cell lines

**Fig. 4A** shows the eight selected human milk-derived peptides and a known bovine milk-derived anticancer peptide, which were synthesized to screen for cancer cytotoxicity by using trypan blue staining. The results showed that HMP-S7, the highest cationic charged peptide (+3 net charge) with α-helical structure, had higher inhibitory activity than the other human milk-derived peptides towards four leukemic cell lines, including Jurkat, Raji, RS4;11, and Sup-B15 cells **(Fig. 4B)**. To study the selectivity of HMP-S7, the effect of HMP-S7 was further observed on normal cells, namely T cells and HEK293T embryonic kidney cells, as well as on solid tumor cell lines, namely SH-SY5Y neuroblastoma, HepG2 hepatoblastoma, A549 lung adenocarcinoma, MDA-MB-231 breast cancer, and HT-29 colorectal adenocarcinoma (**Fig. 4C**). Compared to BMP-S6, a known anti-cancer peptide, HMP-S7 showed less cytotoxic activity towards normal cells and most cancer cells. This suggested that HMP-S7 was more selective to leukemic cells than BMP-S6. Although HMP-S7 showed no cytotoxicity to normal T cells, it had a mild cytotoxic effect on HEK293T cells. Based on these activity profiles, HMP-S7 was chosen to further studies to determine its IC_50_ against four leukemic cell lines (**Fig. 5A**). The results demonstrated the anti-leukemic activity of HMP-S7 was dose-dependent, with the IC_50_ ranging from 89.2 to 186.3 μM depending on the leukemic cell tested. Furthermore, the inhibitory activity of the HMP-S7 treated Jurkat leukemic cell lines was confirmed by the colony-forming assay using soft agar, as shown in **Figure 5B**. Since HMP-S7 could induce leukemic cell death; thereafter, the mechanism of cell death was further investigated.

**Fig. 4.**
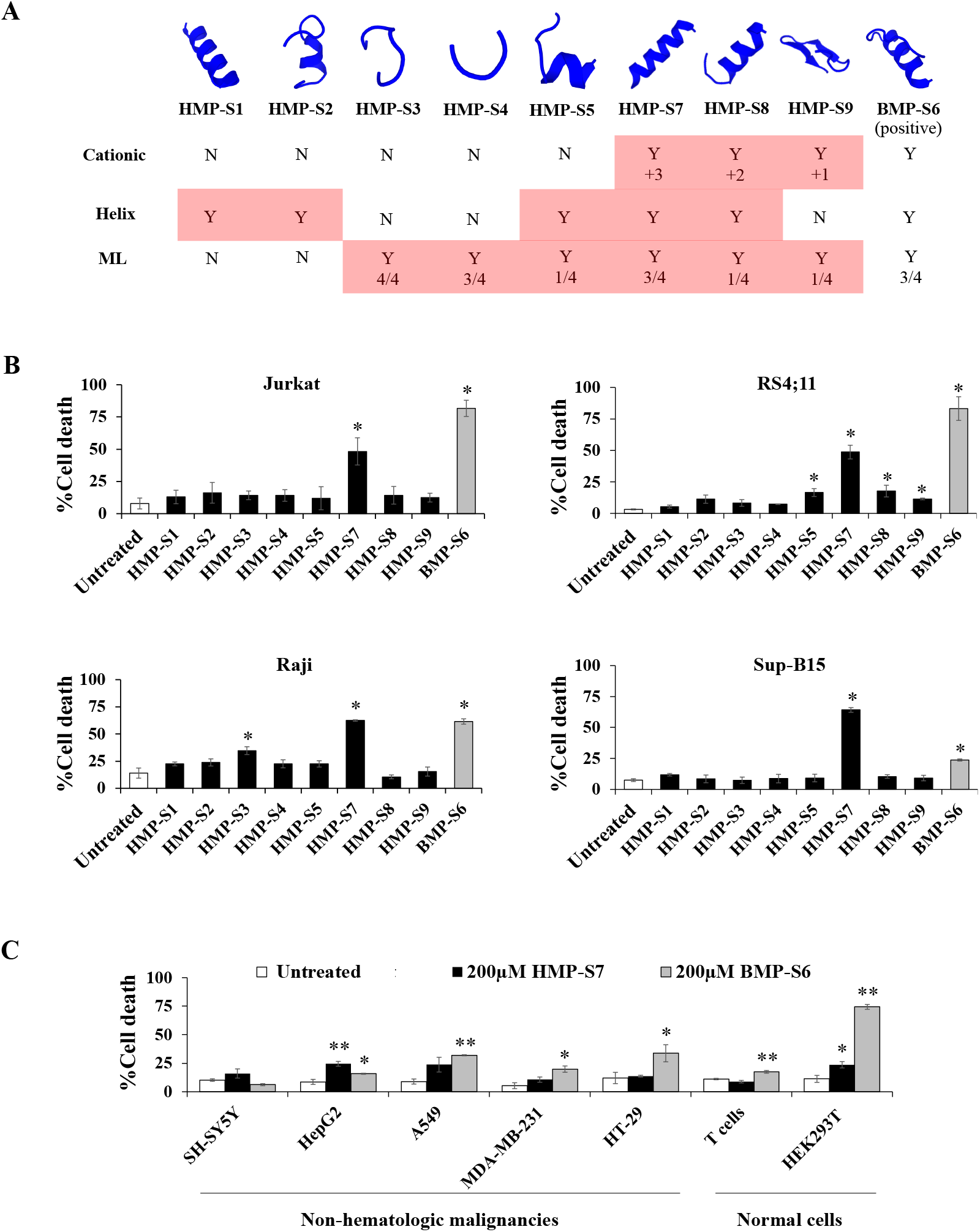
Effect of the 8 synthetic human milk peptides on leukemic and non-leukemic cell lines. (**A**) Eight human milk peptides (HMPs) and a positive control (BMP-S6) were selected and synthesized to test for anti-leukemic activity. The properties of these peptides are summarized, namely predicted secondary structure, cationic nature, helix content, machine learning (ML) prediction of ACP (details of the selected peptides are summarized in **Table S3**). (**B**) Four leukemic cell lines, namely Jurkat, Raji, RS4;11 and Sup-B15, were treated with the 8 synthetic HMPs and the control BMP-S6 at 200 μM, and % cell death was observed after 24 h treatment using the trypan blue exclusion assay under a light microscope. The % cell death was calculated by (number of death cells/total cell number) ×100. The percentage of cell death of all 4 leukemic cell lines after HMP-S7 treatment is significantly increased. (**C**) In addition to leukemic cells, HMP-S7 was also tested on non-hematological malignant cell lines, including neuroblastoma (SH-SY5Y), hepatoblastoma (HepG2), lung cancer (A549), triple-negative breast cancer (MDA-MB-231), colon cancer (HT-29), as well as on normal cells, such as T cells and HEK293T embryonic kidney cells. Statistical significance of differences in % cell death was calculated using three biological replicates, where \**p*<0.05, \*\**p*<0.01 compared to untreated cells.

**Fig. 5.**
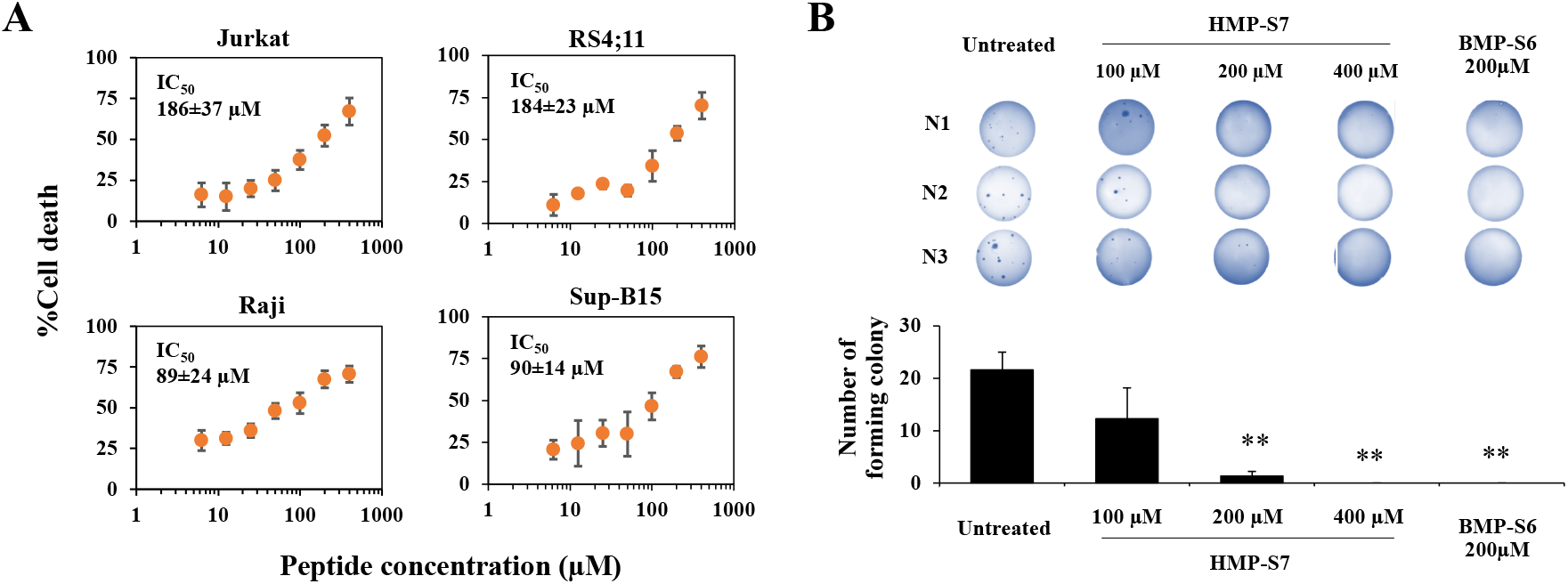
The inhibitory action of HMP-S7 on leukemic cells. (**A**) Four leukemic cell lines, namely Jurkat, Raji, RS4;11, and Sup-B15 cells, were treated with HMP-S7 at various concentrations (0-400 μM) for 24 h and the % cell death (mean ± SD) was determined using trypan blue exclusion assay. (**B**) Effect of HMP-S7 (100 μM, 200 μM, and 400 μM) and the positive control BMP-S6 (200 μM) on colony forming ability of Jurkat cells. After treatment with the test peptide for 24 h, cell suspensions were allowed to form colonies in soft agar for 20 days. The colonies in the soft agar were stained with crystal violet and counted. **, *p*<0.01 compared to untreated condition (n=3 biological replicates).

### Mechanisms of HMP-S7-induced leukemic cell death

To elucidate whether HMP-S7 can act as a membranolytic peptide towards leukemic cells, Jurkat cells were treated with HMP-S7, in untagged form or tagged with fluorescein isothiocyanate (FITC) at the IC_50_ for 24 h. By confocal microscopy, propidium iodide (PI) staining of cytoplasmic RNA was observed in the untagged HMP-7 and the positive control Triton X-100 (**Fig. 6A**). The FITC-tagged HMP-S7 was located in the cytoplasm as with the PI stain of cytoplasmic RNA. Flow cytometry showed that the HMP-S7 could accumulate more in the cytoplasm with increasing concentrations from IC_50_ to 2×IC_50_ (**Fig. 6B** and **6C**). Using the lactate dehydrogenase (LDH) release assay, intracellular LDH leakage into the cell culture supernatant was evident at 24 h in HMP-S7-treated RS4;11 cells (at IC_50_ and 2×IC_50_) and Triton X-100 treated cells (**Fig. 6D**). Finally, HMP-S7-induced leukemic cell death was confirmed by flow cytometry using Annexin V/PI co-staining. The number of apoptotic cells was significantly increased compared to untreated condition, while a dose-response relationship was observed in four leukemic cell lines with IC_50_ and 2×IC_50_ of HMP-S7 treatment (**Fig. 6E** and **6F**). Taken together, our findings suggested that HMP-S7 attacked leukemic cells to form micropores in the cell membranes and penetrated into the cytoplasm, to perturb leukemic cells leading to cell death.

**Fig. 6.**
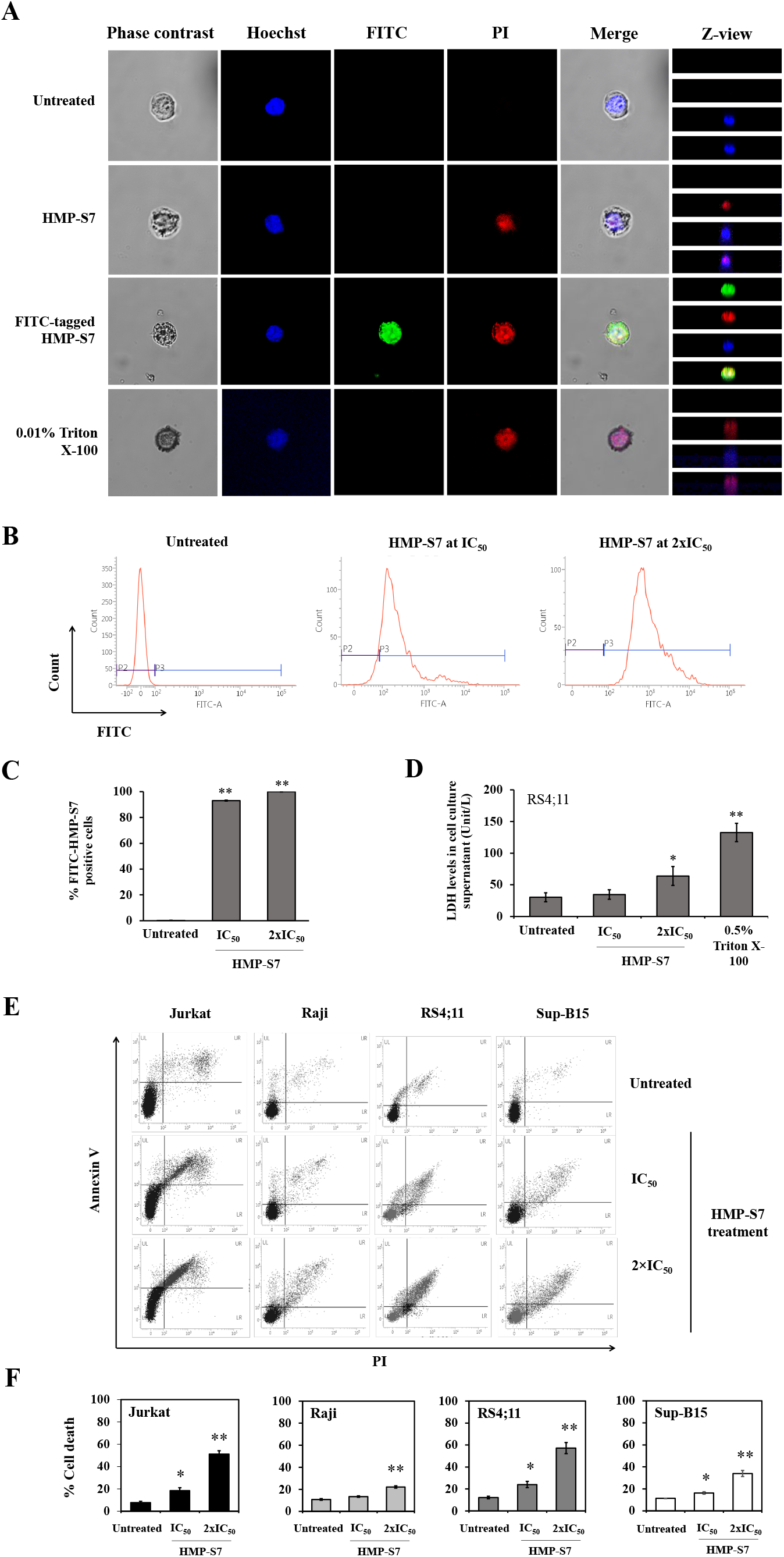
HMP-S7 action on internalization and leukemic cell death. Jurkat cells were treated with HMP-S7 conjugated with or without FITC at IC_50_ and 0.01% Triton X-100 (positive control) and stained with PI to observe membrane permeability **(A)**. The FITC tagged HMP-S7 (at IC_50_ and 2×IC_50_) was internalized into the cytoplasm of Jurkat cells. (**B**) Flow cytometry of FITC tagged HMP-S7-treated Jurkat cells. (**C**) The bar graph showed flow cytometric results as %FITC-positive Jurkat cells. (**D**) LDH release assay showing LDH level in culture supernatant as evidence of cellular membrane disruption of RS4;11 cells treated with HMP-S7 at IC_50_ and 2×IC_50_ for 24 h. Triton X-100 treated cells were used as positive control. (**E**) Flow cytometric cell death assay using Annexin-V/PI co-staining. Four leukemic cell lines, namely Jurkat, Raji, RS4;11, and Sup-B15 were treated with HMP-S7 at IC_50_ and 2×IC_50_ for 24 h. (**F**) Bar graphs showed % cell death composed of upper left (early apoptosis), upper right (late apoptosis) and lower right (necrosis) quadrants of flow cytometric data. \**p*<0.05, \*\**p*<0.01 compared to untreated condition. All experiments were performed in three biological replicates.

### Ex vivo leukemic cytotoxicity of HMP-S7

In the end, we studied whether HMP-S7 can exhibit anti-leukemic effect against patient-derived leukemic cells *ex vivo*. Bone marrow-derived mononuclear cells were isolated from three patients with B cell-acute lymphoblastic leukemia (B-ALL) by Ficoll density gradient centrifugation (demographic data as shown in **Fig. 7A**). The percentage of leukemic cells in bone marrow specimens were 84.5%–92.5% (**Fig. 7A**). Nonetheless, greater than 90% morphologically recognizable malignant lymphoblasts were achieved after mononuclear cell isolation and *ex vivo* culturing (data not shown). These cells were assigned as patient-derived leukemic cells for *ex vivo* leukemic cytotoxic assay using Annexin V/PI co-staining. Interestingly, HMP-S7 (200 μM and 400 μM) exhibited anti-leukemic activity against *ex vivo* leukemic cells isolated from three independent patients in a dose dependent manner (**Fig. 7B** and **7C**). BMP-S6 (200 μM), which demonstrated anticancer activity against multiple cancers including leukemia cell lines (**Fig. 4** and **5**), was included for comparison. However, BMP-S6 had a marginal-to-no cytotoxic effect against patient-derived leukemic cells *ex vivo* (**Fig. 7B** and **7C**). This finding supports further development of HMP-S7 as an anti-leukemic peptide for preclinical and clinical studies.

**Fig. 7.**
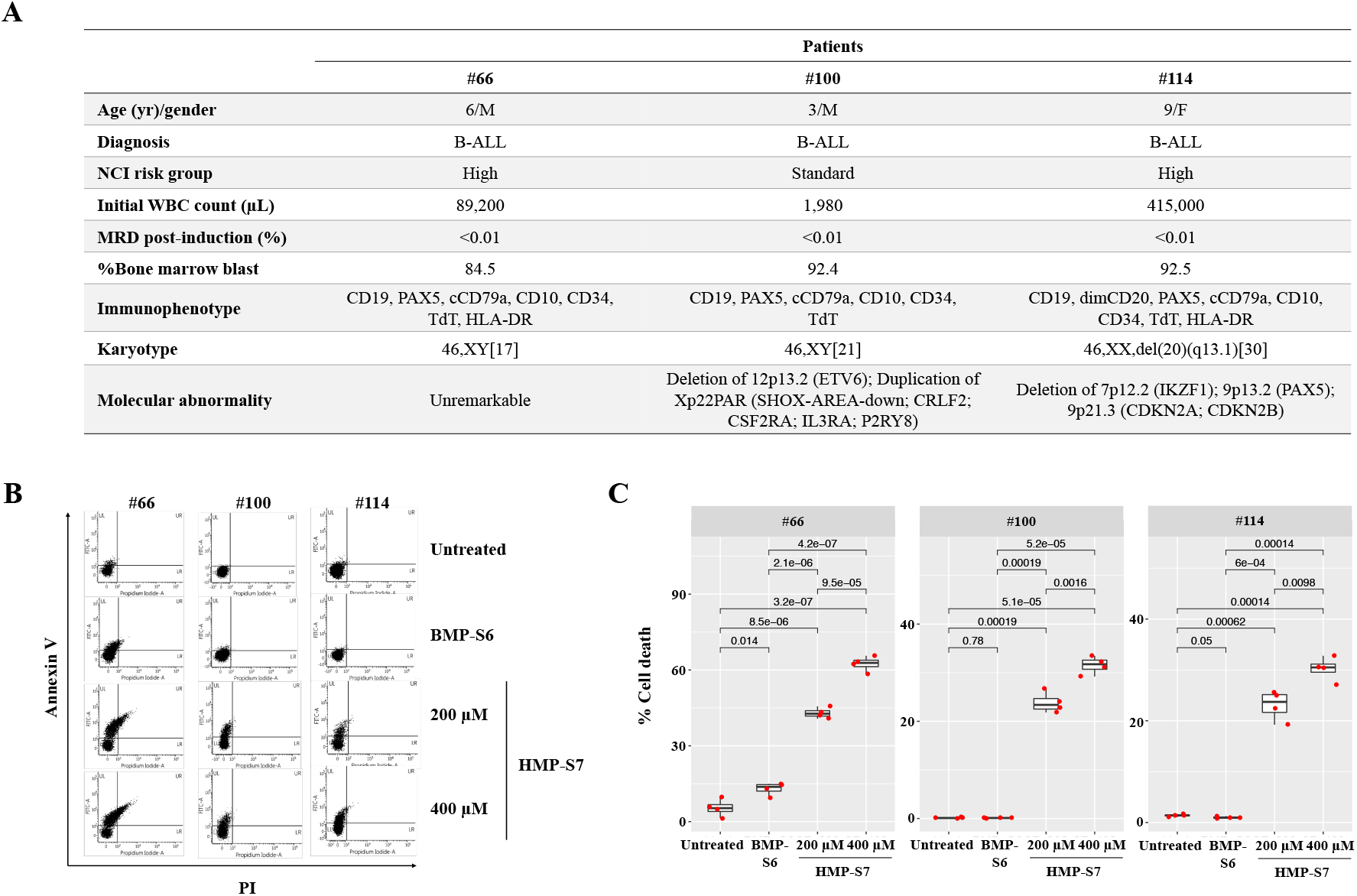
HMP-S7 induced patient-derived leukemic cell death *ex vivo*. Bone marrow derived lymphoblasts were collected from three leukemic patients and were processed as described in “*Materials and Methods*” section. (**A)** Demographic data of three leukemic patients. Patient-derived lymphoblasts were treated with two doses of HMP-S7, and BMP-S6 (a positive control from previous experiments). (**B**) Flow cytometric analysis with annexin V/PI co-staining at 72-h post-treatment. Patient-derived leukemic cells were treated with 200 μM and 400 μM HMP-S7, while 200 μM BMP-S6 was included for comparison. (**C**) Boxplots showed % cell death including upper left, upper right, and lower right quadrants of flow cytometric data (n=4 replicates per condition). Abbreviations: B-ALL, B-cell acute lymphoblastic leukemia; F, female; M, male; MRD, minimal residual disease; NCI, National Cancer Institute; WBC, white blood cells.

### Discussion

This study aimed to search for a novel ACP from naturally occurring human milk peptides. Several reasons suggest that human milk would be a good source for this. First, there is evidence that human breast milk can significantly reduce leukemia in children with more than 6 months of breastfeeding (*25, 26*). Secondly, HAMLET (human alpha-lactalbumin made lethal to tumor cells), a protein-lipid complex that induces apoptosis-like death in tumor cells, was discovered from a human milk protein (*37*). Alpha1H (the alpha1 domain of α-lactalbumin in complex with oleic acid), which was further developed from HAMLET, has now entered the First-in-Human trial in patients with bladder cancer (clinicaltrials.gov identifier NCT03560479) (*38*). Thirdly, ACP has never been explored from human milk. Fourthly, recent advances in mass spectrometric-based proteomics allow high-throughput peptide identification from human milk, and *in silico* ACP prediction algorithms based on peptide sequences have improved in recent years.

Therefore, this study developed an integrative workflow for ACP discovery by combining the strengths of mass spectrometry, *in silico* screening, and experimental validation. This strategy replaces labor-intensive and time-consuming processes of the activity-guided purification by applying mass spectrometric peptide identification of fractionated human milk peptides, and compilation of the results into a peptide library of naturally occurring-human milk peptides. Peptide amino acid sequences were subsequently subjected to *in silico* screening to shorten the time to discovery and lessen the cost of large scale experiments. Finally, the prioritized peptide candidates could be validated for anti-leukemic activities using *in vitro* cellular studies.

To prioritize the peptides for experimental screening and validation, we hypothesized that the consensus results from the physicochemical property (positive charge), secondary peptide structure (alpha-helix), and ACP predictive results from different machine learning models would provide the best chance for identification of ACP from the peptide library. This hypothesis was grounded on previous evidence that; i) the net positively charged peptides are attracted to the net negatively charged cancer cell membranes (*39*); ii) most known oncolytic peptides share alpha-helical structure (*32*); iii) different machine learning models were trained and tested upon various data sets of known ACPs (*33–36*), so these models have varied predictive performance against the new unknown peptide data set. More positive predictions would provide more confidence in the predicted candidates. Since it was unknown at the initial stage of this study whether this integrative approach would be successful, we therefore explored each of the preferred ACP properties. For this reason, eight selected peptide candidates were identified and screened for anti-leukemic effect (**Fig. 3** and **Table S3**). BMP-S6, the positive control of the experimental validation, met all three criteria and showed the anti-leukemic effect with the trade-off of more toxicity toward normal cells. Six out of eight selected peptide candidates did not meet all three criteria and showed no activity *in vitro*. HMP-S7 and HMP-S8 were the top 2 candidates meeting all three criteria of the integrative workflow; nonetheless, only HMP-S7 had cytotoxic effect against leukemic cell lines *in vitro* (**Fig. 4–6**) and patient-derived leukemic cells *ex vivo* (**Fig. 7**). From this observation, we learned that this integrative approach is more accurate than using a single preferred anticancer property to prioritize peptide candidates. When one peptide candidate met all three criteria, the difference in one positive charge (and perhaps two predictions from machine learning models) did matter for the anti-leukemic prediction (**Fig. 3**). In future studies, this integrative approach of ACP screening can be applied to larger data sets, either using mass spectrometric-based or *in silico* generated peptide libraries, to speed up the discovery of ACP against multiple cancer types.

Looking forward, HMP-S7 should be further validated in preclinical models. Further peptide modifications using HMP-S7 as a prototype, for example, amino acid substitutions (*40*), homing motif tagging (*41*), and PEGylation (*42*), may improve anticancer efficacy, tumor targeting, biocompatibility and stability. Furthermore, the human milk peptide library can be expanded by further profiling relevant biological specimens or by *in silico* peptide generation from the proteins of interest. As mentioned for the *in vitro* study support, machine learning for ACP prediction might be further improved by a combination of ensemble models, physicochemical properties, and peptide secondary structure.

This study had limitations. First, it was expected that milk peptides should represent a more significant number of unique identities than those identified in this study. Loss of peptides during mass spectrometric detection due to neutral net charge may be one reason. Some milk peptides may also be lost and degraded during the fractionation process. Addressing these issues would improve the number of unique peptides identified by mass spectrometry in future studies. Secondly, given that the custom-made peptides are commercially available and ready for anticancer activity screening, the issues related to having a small peptide library because of experimental bottlenecks (specimen types, separation processes, instrumental limits of detection) could be addressed by *in silico* generated peptide databases as mentioned above. Thirdly, our workflow is compatible with the native peptides, but this integrative workflow omitted peptides with post-translational modifications (such as glycosylated peptides) by default. Human milk-derived glycopeptidomics could be important for therapeutic peptide investigations, once high-throughput data acquisition strategies and computational predictive tools are tailored for large-scale glycosylated peptide analysis. Lastly, it should be emphasized that *ex vivo* anti-leukemic activity of HMP-S7 was performed on leukemic cells derived from patients with acute lymphoblastic leukemia. Future studies should consider evaluating the anti-leukemic effect of HMP-S7 in a broader context of hematologic malignancies, including acute and chronic myeloid leukemia.

In conclusion, this study applied the integrative workflow to discover HMP-S7 (NH_2_-SFIPRAKSTWLNNIKLL-COOH) as a novel anti-leukemic peptide derived from human breast milk. This anti-leukemic peptide is a cationic peptide with alpha-helical structure that selectively kills leukemic cell lines *in vitro* and exhibits its cytotoxicity against patient-derived leukemic cells *ex vivo*. Future research on peptide modification, together with the efficacy studies in preclinical animal models or early phase clinical trials, is warranted to develop HMP-S7 as peptide-based cancer therapeutics in the future.

### Materials and Methods

#### Experimental design

This study aimed to identify a novel anti-leukemic peptide from human milk. An integrative approach was established by combining the strengths of mass spectrometric-peptide identification for library construction, *in silico* screening of peptide library for prioritizing peptide candidates, and *in vitro* experimental validation for anti-leukemic activities. A newly discovered anti-leukemic peptide was finally examined its activity against patient-derived leukemic cells *ex vivo*. All subjects gave their informed consents for inclusion before they participated in the study. The study was conducted in accordance with the Declaration of Helsinki, and the protocol was approved by the Human Research Ethics Committee, Faculty of Medicine Ramathibodi Hospital, Mahidol University (Protocol ID 11-60-13, No. MURA2017/760; with an approval of amendment on May 3, 2021).

#### Human milk collection

Human milk (60 ml) from 10 healthy volunteer mothers of full-term infant, whose blood test has no infection of HIV, Hepatitis B or C viruses, and Syphilis, were collected by a breast pump and stored in a freezer (−20°C) as per the regulation of Ramathibodi Human Milk Bank (RHMB) before transferring to the laboratory (*43*).

#### Peptide isolation and fractionation

Three pooled milk specimens were produced from 3, 3, and 4 individual specimens (**Table S1**) before further processing. Twenty milliliters of pooled milk specimens were centrifuged at 4°C, 1,500 g for 10 min, and then 5,000 g for 30 min twice to remove cells and lipids. The collected supernatant was centrifuged at 4°C, 12,000 g for 1 h, 32,000 g for 1 h, and finally, 200,000 g for 1 h to remove extracellular vesicles, including microvesicles and exosomes. Thereafter, peptides (<3 kDa) were separated from high molecular size proteins by a 3 kDa cutoff ultrafiltration column (Amicon^®^, Merck Millipore Ltd., Cork, Ireland). The crude peptides (<3 kDa) in the flow-through fraction were injected into the C18 solid-phase extraction (SPE) column (Waters, Waters Corporation, Massachusetts, USA). The bound peptides were then stepwise elution with 1 mL each of 15%, 20%, 25%, 30%, 35%, 40%, 45%, 50%, 55%, and 80% acetonitrile (ACN), respectively. The eluted peptides were dried using a SpeedVac concentrator (Labconco, Labconco Corporation, Missouri, USA). The dried peptides were then resuspended in deionized (d*I*) water for peptide estimation using Bradford’s assay (Bio-rad) (*44*).

#### Synthetic peptides

Nine synthetic peptides (>98% purity) (**Table S3**) were custom ordered from GL Biochem (GL Biochem (Shanghai) Ltd., Shanghai, China) and resuspended in the appropriate culture medium to the final concentration before used.

#### Cell culture

Cell lines were obtained from the American Type Culture Collection (ATCC). Jurkat (ATCC^®^ TIB-152™), RS4;11 (ATCC^®^ CRL-1873™) and Raji (ATCC^®^ CCL-86™) cells were maintained in RPMI-1640 medium (Gibco, Thermo Fisher Scientific, MA, USA) supplemented with 10% fetal bovine serum (FBS) (Gibco) and 1× penicillin/streptomycin (Gibco) in 5% CO_2_ at 37°C, while Sup-B15 (ATCC^®^ CRL-1929™) cells were cultured in Iscove’s modified Dulbecco’s medium (IMDM) supplemented with 0.05 mM 2-mercaptoethanol (Sigma), 20% FBS (Gibco) and 1× penicillin/streptomycin (Gibco). SH-SY5Y (ATCC^®^ CRL-2266™), MDA-MB-231 (ATCC^®^ HTB-26™), A549 (ATCC^®^ CCL-185™) cell lines were grown in Dulbecco’s Modified Eagle’s Medium (DMEM)-high glucose (Gibco) supplemented with 10% FBS and 1× penicillin/streptomycin in 5% CO_2_ at 37°C. HT-29 cells (ATCC^®^ HTB-38™) and HepG2 (ATCC® HB-8065™) were grown in DMEM-F12 (Gibco) supplemented with 10% FBS and 1× penicillin/streptomycin in 5% CO_2_ at 37°C. For representative normal cells, HEK293T human embryonic kidney cells (ATCC^®^ CRL-3216™) were maintained in DMEM-high glucose (Gibco) supplemented with 10% FBS and 1× penicillin/streptomycin in 5% CO_2_ at 37°C. Human peripheral blood mononuclear cells (PBMCs) were prepared from 10 mL EDTA blood for T cell cultivation. The whole blood was diluted in PBS at a ratio of 1:1, and then carefully layered into Ficoll-Paque solution (Robbins Scientific Cooperation, Norway) with a ratio of 2:1 (diluted blood:Ficoll-Paque solution). The tube containing the layered solution was centrifuged at 400 g, 20°C for 35 min with no break. One million PBMCs were cultured in the OKT3 and CD28 coated plate. Briefly, 1 μg each of OKT3 and CD28 was added into 1 ml PBS in a 24-well plate and incubated at room temperature for 2 h. The coated well was washed with PBS once before adding PBMCs. The PBMCs in coated well were cultured in RPMI-1640 supplemented with 100U/mL IL-2, 10% FBS and 1× penicillin/streptomycin in the presence of OKT3 and CD28 for 3 days at 37°C under a humidified atmosphere with 5% CO_2_ before used.

#### Human milk peptide identification by mass spectrometry

Dried human milk peptides (2 μg) were resuspended in 0.1% formic acid and centrifuged at 14,000 rpm for 30 min. The supernatant was collected and injected into the C18 column (75 μm i.d. × 100 mm) by using Easy-nLC (Thermo Fisher Scientific, Inc.) to desalt and concentrate. The peptides were separated with a gradient of 5-45% acetonitrile/0.1% formic acid for 30 min at a flow rate of 300 nl/min. The isolated peptides were identified by the amaZon speed ETD ion trap mass spectrometer (Bruker Daltoniks, Billerica, MA, USA). Peptide sequences and identifications were interpreted using in-house MASCOT software version 2.4.0 with the SwissProt database, against *Homo sapiens*, no fixed and no variable modifications, no enzymatic digestion with no missed cleavage allowed, monoisotopic, ±1.2 Da for peptide tolerance, ±0.6 Da for fragment ion tolerance, 2+ and 3+ charge state for ESI-TRAP instrument. Identified peptides had ion scores higher than 20 (significance threshold at *p* < 0.05).

#### *In silico* anticancer peptide screening

Physicochemical properties of all identified human milk peptides were predicted by using the PepDraw tool (http://www.tulane.edu/~biochem/WW/PepDraw/), including peptide length, mass, net charge, isoelectric point (p*I*), and hydrophobicity. Peptide structures were predicted using the PEP-FOLD3 De novo peptide structure prediction tool (*45–47*). Anticancer properties were predicted by four web-based machine learning programs, including i) ACPred-FL, a sequence base predictor for identifying anti-cancer peptides by using classification mode at confidence 0.5 as default setting (http://server.malab.cn/ACPred-FL/#) (*33*); ii) AntiCP 2.0, an ensemble tree classifier based on amino acid composition (https://webs.iiitd.edu.in/raghava/anticp2/index.html) (*34*); iii) MLACP, a random forest-based prediction of anticancer peptides (https://www.thegleelab.org/MLACP.html) (*35*); and iv) mACPpred, a Support Vector Machine-Based Meta-Predictor (http://www.thegleelab.org/mACPpred/) (*36*).

#### Cytotoxicity assay

For cell treatment, the fractionated milk peptides (10 μg/reaction) or the synthetic peptides (varied concentrations as indicated) were resuspended in 100 μL of culture media with appropriate supplements. For floating cells, the peptide solution was mixed with cell suspension (10,000 cells/10 μL/well) in a well of a 96-well flat-bottomed plate. For adherent cells, 10,000 cells were seeded into a 96-well plate (flat bottom) and cultured until they reached 80% confluence before adding 100 μL of peptide solution into the well containing 80% FHs 74 Int cell confluence (3 technical replication). The peptide-treated cells were incubated for 24, 48, or 72 h in 5% CO_2_ incubator at 37°C with humidity as indicated. Cell viability was measured by trypan blue exclusion or WST-1 assays (Roche Diagnostics GmbH, Mannheim, Germany).

#### Half maximum inhibitory concentration (IC_50_) by trypan blue assay

Four distinct leukemic cell lines, i.e., Jurkat, Raji, RS4;11, and Sup-B15 cell lines were cultured in the medium containing HMP-S7 in vary concentrations (0, 6.25, 12.5, 25, 50, 100, 200, and 400 μM) in 96-well flat-bottomed plates for 24 h (1×10^4^ cells/100 μL/well). The percentage of cell death was estimated using the trypan blue exclusion assay using (cell death number/total cell number) × 100. IC_50_ calculation used linear (y=ax+c) or parabolic (y=ax^2^+bx+c) equation for y=50 value and x value= IC_50_ concentration. IC_50_ concentration of HMP-S7 to the 4 leukemic cell lines was reported as mean±SD.

#### Soft agar assay for colony formation

Base agar (1.5 mL of 0.5% agar containing 1×RPMI supplemented with 10% FBS and 1× penicillin/streptomycin) were plated on each well of a 6-well plate and set aside for 5 min to allow the agar to solidify. Jurkat cells were treated with 100 μM, 200 μM, 400 μM HMP-S7, and 200 μM BMP-S6 in a 96-well plate (3 technical replicates, 100 μL/10,000 cells/well) for 24 h. After 24 h, the treated/untreated cells in the 96-well plate were thoroughly mixed and 20 μL taken to mix in 1 mL of the top agar solution (0.3% agarose containing 1×RPMI supplement with 10% FBS and 1× penicillin/streptomycin). The cell suspension was plated on top of the base agar (3 biological replicates) and then the agarose allowed to solidify. The medium (0.5 mL) was added on top of the agar to prevent the agar from drying. The plate was incubated at 37°C in a humidified incubator for 20 days with medium being added twice a week. The agar was stained with 1 mL of 0.005% crystal violet in 20% ethanol for 1 h, destained with 20% ethanol overnight, and then the colonies were counted under a stereomicroscope SZ61 (Olympus Corporation, Tokyo, Japan).

#### Flow cytometric cell death assay

Four leukemic cell lines (1×10^5^ cells/500 μL/well/cell type) were cultured in 24 well-plate in the medium containing HMP-S7 with IC_50_ and 2×IC_50_ concentrations and untreated condition for 24 h (biological triplicate). The untreated cells were divided into 3 tubes representing unstained cells, cells stained with FITC tagged annexin V (no propidium iodide (PI)), and cells stained with PI (no FITC tagged annexin V) to set up compensation and quadrants. The untreated and treated cells were collected and washed twice with cold PBS. Thereafter, the cells were resuspended with 100 μL of 1× binding buffer and then added 5 μL of FITC tagged annexin V and 5 μL of PI (BD Biosciences, CA, USA). The cell suspensions were gently mixed and incubated at room temperature for 15 min in dark. Then, 400 μL of 1× binding buffer was added to each tube, and then the cells were analyzed by using flow cytometry.

#### Lactate dehydrogenase (LDH) release assay

RS4;11 leukemic cell lines (2×10^5^ cells/1 mL/well/cell type) were cultured in 24 well-plate in the complete medium containing HMP-S7 at IC_50_ and 2×IC_50_ concentrations, without HMP-S7 condition (non-treatment), and 0.5% Triton X-100 for 24 h incubation (biological triplication). The cell suspension was collected and centrifuged at 200 g for 5 min at room temperature. Then 1 mL of supernatant was collected and LDH enzyme was measured from conversion of lactate to pyruvate and NADH, detected as absorbance at 340 nm using Abbott Architect C16000 clinical chemistry analyzer (Holliston, MA, USA) at the Clinical Chemistry Unit, Department of Pathology, Ramathibodi Hospital, Thailand. LDH level (U/L) of the culture media was subtracted with the background of the fresh medium and reported as mean±SD.

#### FITC tagged HMP-S7 internalized into leukemic cells

The 2×10^4^ Jurkat cells/100 μl/well in the 96-well plate were treated with HMP-S7 tagged with/without FITC at IC_50_ for 24 h. Thereafter, the treated and untreated Jurkat cells were washed once with PBS before staining with propidium iodide (PI) cell stain kit (Invitrogen) and Hoechst 33342 (Cell Signaling Technology, Inc, Massachusetts, USA), for 30 min at room temperature in dark. After incubation, the cells were centrifuged at 200 g for 5 min and excess dye was removed before washing twice with PBS. The cell pellet was mixed with 20% glycerol/PBS and mounted on a glass slide for confocal microscopy (Nikon Instruments, Inc, NY, USA).

#### *Ex vivo* leukemic cytotoxicity study

Patient-derived leukemic cells were obtained from Ramathibodi Tumor Biobanking. These cells were isolated from bone marrow aspirated samples obtained from three B-ALL patients by Ficoll-Paque PLUS density gradient centrifugation, washed by sterile PBS, resuspended in FBS/10% DMSO, and then stored in liquid nitrogen until used. Demographic and clinical data were curated from the Electronic Medical Record and patient’s chart. Patient-derived leukemic cells were thawed in a water bath at 37°C for 5 min, washed with sterile PBS, resuspended and maintained *ex vivo* in RPMI supplemented with 20% FBS and 1× penicillin/streptomycin for 7 days. Thereafter, patient-derived leukemic cells were collected by centrifugation at 300 x g for 5 min at room temperature and counted by LUNA-II automated cell counter (Logos Biosystems, Inc., Gyeonggi-do, South Korea). Each specimen was stained by Wright staining and evaluated under a light microscopy to ensure that greater than 90% morphologically recognizable malignant lymphoblasts were detected before further analysis. Approximately 5×10^4^ leukemic cells were then treated with 200 μM BMP-S6, 200 μM and 400 μM HMP-S7 peptides, or not treated (4 replicates per condition) in a 24-well plate. After 3 days of treatment, all cells were collected and washed twice with sterile PBS. The cell pellets were resuspended in 1× binding buffer and then stained with FITC Annexin V/PI apoptosis detection assay (BD Biosciences, CA, USA). The % cell death was determined by using flow cytometry (BD FACSVerse, BD Biosciences).

#### Statistical Analysis

The number and percentage of dead or live cells, OD and IC_50_, were calculated and statistically tested using Student’s *t-test* with statistical significance at *p* < 0.05.

## Acknowledgments

We would like to thank Dr. Natini Jinawath, Principal investigator of Ramathibodi Tumor Biobank, Faculty of Medicine Ramathibodi Hospital, Mahidol University, for providing patient-derived leukemic cells, and Mrs. Sirimon Kongthaworn, for assisting milk collection at RHMB, and staff at the Laboratory of Biochemistry, Chulabhorn Research Institute, and at Pediatric Department and Research Center at Faculty of Medicine Ramathibodi Hospital for providing research instrument facilities.

## Author contributions

Conceptualization: SC; Methodology: WC, SC; Investigation: WC, JP, TV, NP, PP, TS, TL, CS, SC; Visualization: WC, SC; Supervision: JS, SH; Resources, TS, JS, SH, SC; Writing – original draft: WC; Writing – review & editing: JP, TV, NP, PP, TS, TL, CS, JS, SH, SC. All authors have read and agreed to the published version of the manuscript.

## Data and materials availability

All data are available in the main text or the supplementary materials.

