## Supplementary material for "An integrative approach leads to the discovery of a novel anti-leukemic peptide from human milk": Fig. S1; Table S1-S3

### Supplementary figure

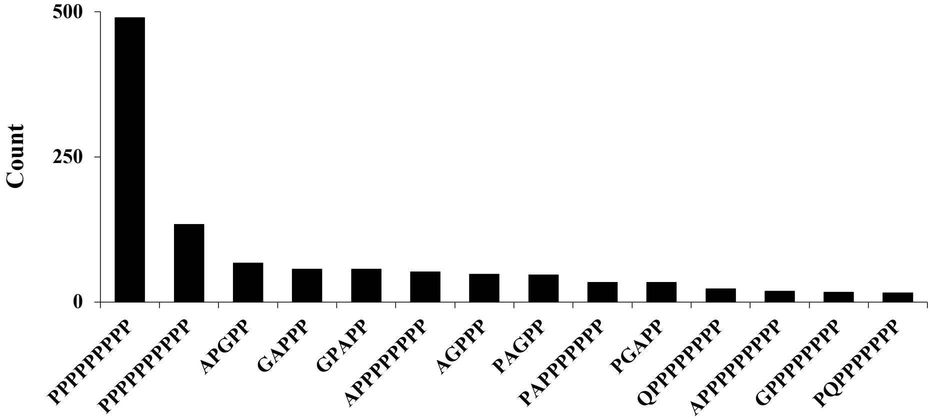


**Fig. S1**. The frequency of the identified peptides with proline-rich sequences.

### Supplementary tables

**Table S1.** Human breast milk data donated by 10 volunteer mothers

| | Volunteer mother no. | Maternal age (year) | Day of lactation (day) |
|---|---|---|---|
| Pooled sample#1 | 1 | 25 | 20 |
| | 2 | 33 | 23 |
| | 3 | 36 | 6 |
| Pooled sample#2 | 4 | 30 | 129 |
| | 5 | 33 | 75 |
| | 6 | 34 | 47 |
| Pooled sample#3 | 7 | 31 | 95 |
| | 8 | 34 | 71 |
| | 9 | 31 | 80 |
| | 10 | 36 | 92 |
| | **Mean** | 32.30 | 63.80 |
| | **SD** | 3.27 | 38.97 |

**Table S2**. An in-house human milk-derived peptide library constructed upon mass spectrometric peptide identification of 11 ACN eluted fractions of naturally occurring human milk peptides with predictions of physicochemical, structural, and anticancer properties.

| Peptide sequence | Structure# | Protein accession | Gene name | Protein name | Protein score | Peptide score | Peptide length | Peptide mass (Da) | Peptide pI | Peptide net charge* | Peptide hydrophobicity (Kcal/mol)* | Extinction coefficient (1/M*1/cm)* | ACN elution fractions | | | | | | | | | | | Frequency | ACPpred-FL | | antiCP 2.0 | | MLACP | | mACPpred | |
|---|---|---|---|---|---|---|---|---|---|---|---|---|---|---|---|---|---|---|---|---|---|---|---|---|---|---|---|---|---|---|---|---|
| | | | | | | | | | | | | | Pooled | 15 % | 20 % | 25 % | 30 % | 35 % | 40 % | 45 % | 50 % | 55 % | 80 % | | Confidence | Prediction | Score | Prediction | Probability | Prediction | Probability | Prediction |
| SFIPRAKSTWLNNIKLL | Helix | O60287 | URB1 | Nucleolar pre-ribosomal-associated protein 1 | 63 | 63.24 | 17 | 2000.15 | 11.70 | 3 | 9.03 | 5500 | | | | | | | | + | | | | 1 | 0.94 | ACP | 0.57 | AntiCP | 0.4010 | Non-ACP | 0.8932 | ACP |
| GRATLVQDGIAKGRVA | Helix-Coil | Q13410 | BTN1A1 | Butyrophilin subfamily 1 member A1 | 65 | 64.69 | 16 | 1610.92 | 11.40 | 2 | 20.64 | 0 | | | | | | | + | + | | | | 2 | 0.83 | ACP | 0.25 | Non-AntiCP | 0.0824 | Non-ACP | 0.2289 | Non-ACP |
| KPPPPPPPP | Coil | A0JLT2 | MED19 | Mediator of RNA polymerase II transcription subunit 19 | 55 | 54.81 | 9 | 922.53 | 10.59 | 1 | 11.82 | 0 | | | | + | | | | | | | | 1 | 0.97 | ACP | 0.58 | AntiCP | 0.6379 | ACP | 0.7355 | ACP |
| PARPPPPPP | Coil | Q9NQC3 | RTN4 | Reticulon-4 | 60 | 59.81 | 9 | 924.52 | 11.56 | 1 | 11.19 | 0 | + | | | | | | | | | | | 1 | 0.99 | ACP | 0.68 | AntiCP | 0.6492 | ACP | 0.9665 | ACP |
| VPKAKDTVYT | Helix-Coil | P05814 | CSN2 | Beta-casein | 293 | 60.05 | 10 | 1120.61 | 9.42 | 1 | 16.65 | 1490 | | | | + | | | | | | | | 1 | 0.99 | ACP | 0.29 | Non-AntiCP | 0.4242 | Non-ACP | 0.5130 | Non-ACP |
| LWSVPQPKVLPIPQ | Coil | P05814 | CSN2 | Beta-casein | 231 | 70.83 | 14 | 1600.93 | 9.84 | 1 | 6.63 | 5500 | | | | + | | + | | + | | | | 3 | 0.99 | ACP | 0.3 | Non-AntiCP | 0.4217 | Non-ACP | 0.7948 | ACP |
| QVVPYPQRAVPVQA | Sheet-Coil-Sheet | P05814 | CSN2 | Beta-casein | 293 | 42.56 | 14 | 1550.85 | 9.97 | 1 | 10.89 | 1490 | | | | + | | | | | | | | 1 | 0.99 | ACP | 0.2 | Non-AntiCP | 0.5000 | Non-ACP | 0.1697 | Non-ACP |
| LALPPQPLWSVPQPK | Coil | P05814 | CSN2 | Beta-casein | 231 | 57.87 | 15 | 1669.95 | 9.80 | 1 | 7.60 | 5500 | | | | | | | | + | | | | 1 | 0.99 | ACP | 0.34 | Non-AntiCP | 0.4070 | Non-ACP | 0.7903 | ACP |
| LPIPQQVVPYPQRAVPVQ | Sheet-Coil-Sheet | P05814 | CSN2 | Beta-casein | 328 | 56.28 | 18 | 2028.15 | 9.60 | 1 | 9.07 | 1490 | | | | | | + | | | | | | 1 | 0.99 | ACP | 0.33 | Non-AntiCP | 0.4242 | Non-ACP | 0.0783 | Non-ACP |
| PPQPLWSVPQPKVLPIPQ | Coil | P05814 | CSN2 | Beta-casein | 213 | 55.56 | 18 | 2020.15 | 9.84 | 1 | 7.82 | 5500 | | | | | | | + | | | | | 1 | 0.99 | ACP | 0.25 | Non-AntiCP | 0.3833 | Non-ACP | 0.6944 | ACP |
| AGPPP | Coil | Q96IF1 | AJUBA | LIM domain-containing protein ajuba | 20 | 20.33 | 5 | 437.23 | 5.65 | 0 | 9.97 | 0 | | | | + | | | | | | | | 1 | NA | NP | 0.6 | AntiCP | 0.5883 | ACP | 0.9273 | ACP |
| APGPP | Coil | Q96P50 | ACAP3 | Arf-GAP with coiled-coil, ANK repeat and PH domain-containing protein 3 | 20 | 20.33 | 5 | 437.23 | 5.65 | 0 | 9.97 | 0 | | | | + | | | | | | | | 1 | NA | NP | 0.6 | AntiCP | 0.5979 | ACP | 0.9276 | ACP |
| GAPPP | Coil | P35368 | ADRA1B | Alpha-1B adrenergic receptor | 20 | 20.33 | 5 | 437.23 | 5.65 | 0 | 9.97 | 0 | | | | + | | | | | | | | 1 | NA | NP | 0.6 | AntiCP | 0.5734 | ACP | 0.9750 | ACP |
| GPAPP | Coil | Q99490 | AGAP2 | Arf-GAP with GTPase, ANK repeat and PH domain-containing protein 2 | 20 | 20.33 | 5 | 437.23 | 5.65 | 0 | 9.97 | 0 | | | | + | | | | | | | | 1 | NA | NP | 0.6 | AntiCP | 0.5904 | ACP | 0.9768 | ACP |
| PAGPP | Coil | O43306 | ADCY6 | Adenylate cyclase type 6 | 20 | 20.33 | 5 | 437.23 | 5.25 | 0 | 9.97 | 0 | | | | + | | | | | | | | 1 | NA | NP | 0.6 | AntiCP | 0.6000 | ACP | 0.9689 | ACP |
| PGAPP | Coil | O15013 | ARHGEF10 | Rho guanine nucleotide exchange factor 10 | 20 | 20.33 | 5 | 437.23 | 5.25 | 0 | 9.97 | 0 | | | | + | | | | | | | | 1 | NA | NP | 0.6 | AntiCP | 0.5940 | ACP | 0.9284 | ACP |
| AIQDPRLF | Coil | P01833 | PIGR | Polymeric immunoglobulin receptor | 93 | 49.14 | 8 | 958.52 | 6.56 | 0 | 10.68 | 0 | | | | + | | | | | | | | 1 | 0.61 | Non-ACP | 0.34 | Non-AntiCP | 0.4447 | Non-ACP | 0.3963 | Non-ACP |
| LPNSHPPT | Coil | P07498 | CSN3 | Kappa-casein | 60 | 45.08 | 8 | 861.43 | 7.57 | 0 | 10.96 | 0 | | | | + | | | | | | | | 1 | 0.97 | ACP | 0.34 | Non-AntiCP | 0.3629 | Non-ACP | 0.8704 | ACP |
| YQPPPPPP | Coil | P49715 | CEBPA | CCAAT/enhancer-binding protein alpha | 74 | 58.72 | 8 | 891.45 | 5.48 | 0 | 8.80 | 1490 | + | + | + | | | | | | | | | 3 | 0.94 | ACP | 0.41 | Non-AntiCP | 0.6250 | ACP | 0.8007 | ACP |
| APPPPPPPP | Coil | P46379 | BAG6 | Large proline-rich protein BAG6 | 121 | 65.46 | 9 | 865.47 | 5.65 | 0 | 9.52 | 0 | + | + | + | | | | | | | | | 3 | 0.86 | Non-ACP | 0.58 | AntiCP | 0.6373 | ACP | 0.9284 | ACP |

| | | | | | | | | | | | | | | | | | | | | | | | | | | |
|---|---|---|---|---|---|---|---|---|---|---|---|---|---|---|---|---|---|---|---|---|---|---|---|---|---|---|
| GHPPPPPPP | Coil | P14866 | HNRNPL | Heterogeneous nuclear ribonucleoprotein L | 107 | 76.63 | 9 | 891.46 | 8.32 | 0 | 12.36 | 0 | + | + | + | + | | 4 | 0.98 | ACP | 0.76 | AntiCP | 0.6166 | ACP | 0.9766 | ACP |
| GPPPPPPPP | Coil | P10275 | AR | Androgen receptor | 64 | 64.05 | 9 | 851.45 | 5.65 | 0 | 10.17 | 0 | + | + | | | | 2 | 0.64 | ACP | 0.84 | AntiCP | 0.6374 | ACP | 0.9766 | ACP |
| HPPPPPPPP | Coil | Q9C0F0 | ASXL3 | Putative Polycomb group protein ASXL3 | 66 | 42.03 | 9 | 931.49 | 8.32 | 0 | 11.35 | 0 | | | | + | | 1 | 0.94 | ACP | 0.63 | AntiCP | 0.6350 | ACP | 0.9281 | ACP |
| LPLPPPPPP | Coil | Q5VT03 | NUTM2D | NUT family member 2D | 62 | 61.59 | 9 | 923.55 | 5.63 | 0 | 6.38 | 0 | | | | + | | 1 | 0.99 | ACP | 0.54 | AntiCP | 0.6337 | ACP | 0.9288 | ACP |
| LPNSHPPTV | Coil | P07498 | CSN3 | Kappa-casein | 60 | 61.74 | 9 | 960.50 | 7.89 | 0 | 10.50 | 0 | | | | + | | 1 | 0.99 | ACP | 0.34 | Non-AntiCP | 0.2683 | Non-ACP | 0.7205 | ACP |
| LPPLPPPPP | Coil | P51608 | MECP2 | Methyl-CpG-binding protein 2 | 69 | 69.14 | 9 | 923.55 | 5.63 | 0 | 6.38 | 0 | | | | + | + | 2 | 0.97 | ACP | 0.54 | AntiCP | 0.6337 | ACP | 0.9286 | ACP |
| LPPPLPPPP | Coil | Q86VE0 | MYPOP | Myb-related transcription factor, partner of profilin | 78 | 62.75 | 9 | 923.55 | 5.63 | 0 | 6.38 | 0 | | | | + | | 1 | 0.82 | ACP | 0.54 | AntiCP | 0.6337 | ACP | 0.9785 | ACP |
| PAPPPPPPP | Coil | Q12830 | BPTF | Nucleosome-remodeling factor subunit BPTF | 103 | 81.84 | 9 | 865.47 | 5.25 | 0 | 9.52 | 0 | + | + | + | | | 3 | 0.97 | ACP | 0.58 | AntiCP | 0.6350 | ACP | 0.9686 | ACP |
| PGPPPPPPP | Coil | Q2V2M9 | FHOD3 | FH1/FH2 domain-containing protein 3 | 64 | 64.05 | 9 | 851.45 | 5.25 | 0 | 10.17 | 0 | + | + | | | | 2 | 0.82 | ACP | 0.84 | AntiCP | 0.6363 | ACP | 0.9284 | ACP |
| PHPPPPPPP | Coil | Q15390 | MTFR1 | Mitochondrial fission regulator 1 | 42 | 42.03 | 9 | 931.49 | 8.32 | 0 | 11.35 | 0 | | | | + | | 1 | 0.99 | ACP | 0.63 | AntiCP | 0.6304 | ACP | 0.9289 | ACP |
| PLLPPPPPP | Coil | O95450 | ADAMTS2 | A disintegrin and metalloproteinase with thrombospondin motifs 2 | 80 | 76.11 | 9 | 923.55 | 5.25 | 0 | 6.38 | 0 | + | | | + | | 2 | 0.99 | ACP | 0.54 | AntiCP | 0.6321 | ACP | 0.9441 | ACP |
| PLPIPPPPP | Coil | Q8IX15 | HOMEZ | Homeobox and leucine zipper protein Homez | 73 | 72.62 | 9 | 923.55 | 5.25 | 0 | 6.51 | 0 | | | | + | | 1 | 0.94 | ACP | 0.56 | AntiCP | 0.6163 | ACP | 0.9356 | ACP |
| PLPLPPPPP | Coil | Q13495 | MAMLD1 | Mastermind-like domain-containing protein 1 | 73 | 72.62 | 9 | 923.55 | 5.25 | 0 | 6.38 | 0 | | | | + | | 1 | 0.99 | ACP | 0.54 | AntiCP | 0.6288 | ACP | 0.9357 | ACP |
| PLPPLPPPP | Coil | Q8NEA6 | GLIS3 | Zinc finger protein GLIS3 | 66 | 65.65 | 9 | 923.55 | 5.25 | 0 | 6.38 | 0 | | | | + | | 1 | 0.94 | ACP | 0.54 | AntiCP | 0.6337 | ACP | 0.9783 | ACP |
| PPAPPPPPP | Coil | Q8NFC6 | BOD1L1 | Biorientation of chromosomes in cell division protein 1-like 1 | 51 | 57.78 | 9 | 865.47 | 5.25 | 0 | 9.52 | 0 | | + | | | | 1 | 0.97 | ACP | 0.58 | AntiCP | 0.6369 | ACP | 0.9286 | ACP |
| PPGPPPPPP | Coil | P0C7U0 | ELFN1 | Protein ELFN1 | 56 | 56.49 | 9 | 851.45 | 5.25 | 0 | 10.17 | 0 | + | | | | | 1 | 0.97 | ACP | 0.84 | AntiCP | 0.6377 | ACP | 0.9285 | ACP |
| PPIIPPPPP | Coil | Q16637 | SMN1 | Survival motor neuron protein | 91 | 70.99 | 9 | 923.55 | 5.25 | 0 | 6.64 | 0 | | | | + | | 1 | 0.99 | ACP | 0.6 | AntiCP | 0.6254 | ACP | 0.9451 | ACP |
| PPLLPPPPP | Coil | O95450 | ADAMTS2 | A disintegrin and metalloproteinase with thrombospondin motifs 2 | 80 | 58.31 | 9 | 923.55 | 5.25 | 0 | 6.38 | 0 | | | | + | | 1 | 0.99 | ACP | 0.54 | AntiCP | 0.6321 | ACP | 0.9375 | ACP |
| PPPGSFPPP | Coil | Q15427 | SF3B4 | Splicing factor 3B subunit 4 | 62 | 62.44 | 9 | 891.45 | 5.25 | 0 | 8.64 | 0 | | + | | | | 1 | 0.94 | ACP | 0.71 | AntiCP | 0.6075 | ACP | 0.9103 | ACP |
| PPPPPPPHG | Coil | Q16676 | FOXD1 | Forkhead box protein D1 | 66 | 64.33 | 9 | 891.46 | 7.91 | 0 | 12.36 | 0 | + | + | + | + | | 4 | 0.82 | ACP | 0.76 | AntiCP | 0.6173 | ACP | 0.9276 | ACP |
| PPPPPPPPP | Coil | P10323 | ACR | Acrosin | 155 | 84.84 | 9 | 891.48 | 5.25 | 0 | 9.16 | 0 | + | + | + | + | | 4 | 0.82 | ACP | 0.67 | AntiCP | 0.6377 | ACP | 0.9287 | ACP |
| PQPPPPPPP | Coil | Q6UUV9 | CRTC1 | CREB-regulated transcription coactivator 1 | 56 | 56.40 | 9 | 922.49 | 5.25 | 0 | 9.79 | 0 | | | + | | | 1 | 0.94 | ACP | 0.46 | AntiCP | 0.6350 | ACP | 0.8072 | ACP |

| | | | | | | | | | | | | | | | | | | | | | | | | | | | | | | | | |
|---|---|---|---|---|---|---|---|---|---|---|---|---|---|---|---|---|---|---|---|---|---|---|---|---|---|---|---|---|---|---|---|---|
| PVQPPPPPP | Coil | Q16204 | CCDC6 | Coiled-coil domain-containing protein 6 | 114 | 62.54 | 9 | 924.51 | 5.25 | 0 | 9.19 | 0 | + | | | | | | | | | | | 1 | 0.99 | ACP | 0.27 | Non-AntiCP | 0.6175 | ACP | 0.7744 | ACP |
| QPPPPPPPP | Coil | O14514 | BAI1 | Brain-specific angiogenesis inhibitor 1 | 71 | 54.81 | 9 | 922.49 | 5.47 | 0 | 9.79 | 0 | | | | + | | | | | | | | 1 | 0.94 | ACP | 0.46 | AntiCP | 0.6373 | ACP | 0.7691 | ACP |
| QYPPPPPPP | Coil | Q92841 | DDX17 | Probable ATP-dependent RNA helicase DDX17 | 83 | 67.97 | 9 | 988.50 | 5.45 | 0 | 8.94 | 1490 | + | + | | + | | | | | | | | 3 | 0.94 | ACP | 0.46 | AntiCP | 0.6258 | ACP | 0.9093 | ACP |
| SFPPPPPPP | Coil | B1AK53 | ESPN | Espin | 66 | 42.03 | 9 | 931.48 | 5.49 | 0 | 7.63 | 0 | | | | + | | | | | | | | 1 | 0.94 | ACP | 0.66 | AntiCP | 0.6428 | ACP | 0.9087 | ACP |
| YAPPPPPPP | Coil | Q75N03 | CBLL1 | E3 ubiquitin-protein ligase Hakai | 42 | 42.03 | 9 | 931.48 | 5.48 | 0 | 8.67 | 1490 | | | | + | | | | | | | | 1 | 0.97 | ACP | 0.62 | AntiCP | 0.6253 | ACP | 0.9682 | ACP |
| YQPPPPPPP | Coil | P49715 | CEBPA | CCAAT/enhancer-binding protein alpha | 74 | 57.49 | 9 | 988.50 | 5.48 | 0 | 8.94 | 1490 | + | + | | | | | | | | | | 2 | 0.94 | ACP | 0.46 | AntiCP | 0.6240 | ACP | 0.8092 | ACP |
| APPPPPPPPP | Coil | P46379 | BAG6 | Large proline-rich protein BAG6 | 209 | 55.20 | 10 | 962.52 | 5.65 | 0 | 9.66 | 0 | | + | + | | | | | | | | | 2 | 0.86 | Non-ACP | 0.58 | AntiCP | 0.6372 | ACP | 0.9285 | ACP |
| GHPPPPPPPP | Coil | P14866 | HNRNPL | Heterogeneous nuclear ribonucleoprotein L | 154 | 66.03 | 10 | 988.51 | 8.32 | 0 | 12.50 | 0 | + | + | + | | | | | | | | | 3 | 0.93 | ACP | 0.75 | AntiCP | 0.6268 | ACP | 0.9766 | ACP |
| PAPPPPPPPP | Coil | Q12830 | BPTF | Nucleosome-remodeling factor subunit BPTF | 103 | 55.20 | 10 | 962.52 | 5.25 | 0 | 9.66 | 0 | | + | + | | | | | | | | | 2 | 0.97 | ACP | 0.58 | AntiCP | 0.6368 | ACP | 0.9687 | ACP |
| PPPPPPPPPP | Coil | Q5TGY3 | AHDC1 | AT-hook DNA-binding motif-containing protein 1 | 177 | 66.03 | 10 | 988.54 | 5.25 | 0 | 9.30 | 0 | + | + | + | | | | | | | | | 3 | 0.82 | ACP | 0.67 | AntiCP | 0.6373 | ACP | 0.9287 | ACP |
| SPAVLVHRDG | Helix | Q13410 | BTN1A1 | Butyrophilin subfamily 1 member A1 | 51 | 50.51 | 10 | 1049.56 | 7.91 | 0 | 15.76 | 0 | | | | + | | | | | | | | 1 | 0.97 | ACP | 0.22 | Non-AntiCP | 0.3919 | Non-ACP | 0.4451 | Non-ACP |
| DKIYPSFQPQP | Helix-Coil | P05814 | CSN2 | Beta-casein | 293 | 68.27 | 11 | 1318.65 | 6.79 | 0 | 13.22 | 1490 | | | | + | | | | | | | | 1 | 0.61 | Non-ACP | 0.4 | Non-AntiCP | 0.3709 | Non-ACP | 0.1729 | Non-ACP |
| LPIIQKLEPQI | Helix-Coil | Q99541 | PLIN2 | Perilipin-2 | 49 | 44.53 | 11 | 1290.79 | 6.82 | 0 | 10.29 | 0 | | | | + | | | | | | | | 1 | 0.99 | ACP | 0.27 | Non-AntiCP | 0.4362 | Non-ACP | 0.4128 | Non-ACP |
| PLAPVHNPISV | Coil | P05814 | CSN2 | Beta-casein | 178 | 67.84 | 11 | 1142.64 | 7.89 | 0 | 9.17 | 0 | | | | + | | + | | | + | + | | 4 | 0.99 | ACP | 0.44 | Non-AntiCP | 0.4213 | Non-ACP | 0.5990 | ACP |
| EVSRLQGTGGPS | Helix-Coil | O75888 | TNFSF13 | Tumor necrosis factor ligand superfamily member 13 | 46 | 45.65 | 12 | 1186.59 | 6.61 | 0 | 17.16 | 0 | | | | + | | | | | | | | 1 | 0.61 | Non-ACP | 0.27 | Non-AntiCP | 0.2058 | Non-ACP | 0.0548 | Non-ACP |
| LPIIQKLEPQIA | Helix-Coil | Q99541 | PLIN2 | Perilipin-2 | 49 | 39.04 | 12 | 1361.83 | 6.86 | 0 | 10.79 | 0 | | | | + | + | | | | | | | 2 | 0.99 | ACP | 0.27 | Non-AntiCP | 0.3919 | Non-ACP | 0.4496 | Non-ACP |
| VSLISAEVPLGR | Helix-Coil | Q6PP77 | XKRX | XK-related protein 2 | 53 | 53.14 | 12 | 1239.72 | 6.51 | 0 | 11.51 | 0 | | | | | | | | | | | + | 1 | 0.99 | ACP | 0.14 | Non-AntiCP | 0.3000 | Non-ACP | 0.0435 | Non-ACP |
| TQPLAPVHNPISV | Sheet-Coil-Sheet | P05814 | CSN2 | Beta-casein | 293 | 41.70 | 13 | 1371.75 | 7.89 | 0 | 10.19 | 0 | | | | + | | | | | | | | 1 | 0.98 | ACP | 0.21 | Non-AntiCP | 0.2486 | Non-ACP | 0.2319 | Non-ACP |
| YEKVSAGNGGSSL | Helix-Coil | P15941 | MUC1 | Mucin-1 | 374 | 64.27 | 13 | 1267.60 | 6.75 | 0 | 18.09 | 1490 | + | + | + | + | | | | | | | | 4 | 0.99 | ACP | 0.23 | Non-AntiCP | 0.2421 | Non-ACP | 0.0490 | Non-ACP |
| SPYEKVSAGNGGSS | Helix-Coil | P15941 | MUC1 | Mucin-1 | 245 | 61.61 | 14 | 1338.60 | 6.57 | 0 | 19.94 | 1490 | | | | + | | | | | | | | 1 | 0.99 | ACP | 0.29 | Non-AntiCP | 0.3666 | Non-ACP | 0.0394 | Non-ACP |
| LLQPLMQQVPQPIPQ | Helix-Coil | P05814 | CSN2 | Beta-casein | 328 | 58.79 | 15 | 1728.96 | 5.48 | 0 | 6.31 | 0 | | | | + | + | | | | | | | 2 | 0.99 | ACP | 0.19 | Non-AntiCP | 0.3854 | Non-ACP | 0.2150 | Non-ACP |
| PVTQPLAPVHNPISV | Coil | P05814 | CSN2 | Beta-casein | 293 | 39.05 | 15 | 1567.87 | 7.89 | 0 | 9.87 | 0 | | | + | + | | + | | | | | | 3 | 0.98 | ACP | 0.23 | Non-AntiCP | 0.3877 | Non-ACP | 0.3339 | Non-ACP |
| SPYEKVSAGNGGSSL | Helix-Coil | P15941 | MUC1 | Mucin-1 | 245 | 77.25 | 15 | 1451.69 | 6.75 | 0 | 18.69 | 1490 | + | | + | + | | | | | | | | 3 | 0.99 | ACP | 0.25 | Non-AntiCP | 0.2774 | Non-ACP | 0.0377 | Non-ACP |
| YEKVSAGNGGSSLSY | Sheet-Coil-Sheet | P15941 | MUC1 | Mucin-1 | 374 | 70.58 | 15 | 1517.70 | 6.55 | 0 | 17.84 | 2980 | | | + | | | | | | | | | 1 | 0.99 | ACP | 0.2 | Non-AntiCP | 0.3305 | Non-ACP | 0.0349 | Non-ACP |
| LLQPLMQQVPQPIPQT | Helix-Coil | P05814 | CSN2 | Beta-casein | 293 | 60.08 | 16 | 1830.00 | 5.47 | 0 | 6.56 | 0 | | | | + | | | | | | | | 1 | 0.99 | ACP | 0.17 | Non-AntiCP | 0.3114 | Non-ACP | 0.0613 | Non-ACP |
| LPGQGVHSQGQGPGAN | Coil | Q92896 | GLG1 | Golgi apparatus protein 1 | 49 | 49.37 | 16 | 1502.72 | 7.38 | 0 | 18.67 | 0 | | | | + | | | | | | | | 1 | 0.98 | ACP | 0.23 | Non-AntiCP | 0.2935 | Non-ACP | 0.0411 | Non-ACP |
| SPYEKVSAGNGGSSLS | Helix-Coil | P15941 | MUC1 | Mucin-1 | 191 | 88.19 | 16 | 1538.72 | 6.57 | 0 | 19.15 | 1490 | + | | + | + | | | | | | | | 3 | 0.99 | ACP | 0.28 | Non-AntiCP | 0.3215 | Non-ACP | 0.0456 | Non-ACP |

| | | | | | | | | | | | | | | | | | | | | | | | | | | | | | |
|---|---|---|---|---|---|---|---|---|---|---|---|---|---|---|---|---|---|---|---|---|---|---|---|---|---|---|---|---|---|
| DRSPYEKVSAGNGGSSL | Coil-Helix-Coil | P15941 | MUC1 | Mucin-1 | 245 | 51.74 | 17 | 1722.81 | 6.87 | 0 | 24.14 | 1490 | | | | + | | | | | 1 | 0.94 | Non-ACP | 0.15 | Non-AntiCP | 0.1445 | Non-ACP | 0.0291 | Non-ACP |
| IYPVTQPLAPVHNPISV | Coil | P05814 | CSN2 | Beta-casein | 328 | 55.46 | 17 | 1844.02 | 7.80 | 0 | 8.04 | 1490 | | | | | | + | | | 1 | 0.87 | ACP | 0.21 | Non-AntiCP | 0.2986 | Non-ACP | 0.1128 | Non-ACP |
| SPYEKVSAGNGGSSLSY | Helix-Coil | P15941 | MUC1 | Mucin-1 | 374 | 91.21 | 17 | 1701.78 | 6.55 | 0 | 18.44 | 2980 | | | + | + | | | | | 2 | 0.99 | ACP | 0.25 | Non-AntiCP | 0.3478 | Non-ACP | 0.0382 | Non-ACP |
| DRSPYEKVSAGNGGSSLS | Coil-Helix-Coil | P15941 | MUC1 | Mucin-1 | 245 | 54.92 | 18 | 1809.85 | 6.69 | 0 | 24.60 | 1490 | | | | + | | | | | 1 | 0.86 | Non-ACP | 0.14 | Non-AntiCP | 0.1844 | Non-ACP | 0.0308 | Non-ACP |
| DRSPYEKVSAGNGGSSLSY | Helix-Coil | P15941 | MUC1 | Mucin-1 | 245 | 71.44 | 19 | 1972.91 | 6.67 | 0 | 23.89 | 2980 | | | + | + | | | | | 2 | 0.94 | Non-ACP | 0.12 | Non-AntiCP | 0.2148 | Non-ACP | 0.0291 | Non-ACP |
| TDRSPYEKVSAGNGGSSLS | Helix-Coil | P15941 | MUC1 | Mucin-1 | 374 | 74.97 | 19 | 1910.89 | 6.69 | 0 | 24.85 | 1490 | | | + | + | | | | | 2 | 0.98 | ACP | 0.13 | Non-AntiCP | 0.1328 | Non-ACP | 0.0273 | Non-ACP |
| TDRSPYEKVSAGNGGSSLSY | Coil-Helix-Coil | P15941 | MUC1 | Mucin-1 | 245 | 37.86 | 20 | 2073.96 | 6.67 | 0 | 24.14 | 2980 | | | | + | | | | | 1 | 0.98 | ACP | 0.11 | Non-AntiCP | 0.1655 | Non-ACP | 0.0296 | Non-ACP |
| PTHQIYPVTQPLAPVHNPISV | Sheet-Coil | P05814 | CSN2 | Beta-casein | 293 | 57.77 | 21 | 2307.23 | 7.96 | 0 | 11.53 | 1490 | | | | + | | | | | 1 | 0.98 | ACP | 0.28 | Non-AntiCP | 0.2176 | Non-ACP | 0.0613 | Non-ACP |
| TDRSPYEKVSAGNGGSSLSY TNPAVAATSANL | Coil | P15941 | MUC1 | Mucin-1 | 245 | 66.88 | 32 | 3184.52 | 6.82 | 0 | 27.23 | 2980 | | | | + | | | | | 1 | 0.93 | ACP | 0.12 | Non-AntiCP | 0.0538 | Non-ACP | 0.0326 | Non-ACP |
| PEPPPPPPP | Coil | Q15911 | ZFHX3 | Zinc finger homeobox protein 3 | 81 | 65.88 | 9 | 923.47 | 3.01 | -1 | 12.65 | 0 | + | | | + | + | | | | 3 | 0.94 | ACP | 0.71 | AntiCP | 0.6358 | ACP | 0.7755 | ACP |
| DAPPPPAAPLP | Coil | Q99523 | SORT1 | Sortilin | 76 | 70.38 | 11 | 1041.55 | 2.95 | -1 | 12.63 | 0 | | + | | | | | | | 1 | 0.86 | Non-ACP | 0.37 | Non-AntiCP | 0.5191 | ACP | 0.9143 | ACP |
| PQIPKLTDLEN | Helix-Coil | P05814 | CSN2 | Beta-casein | 231 | 67.85 | 11 | 1266.68 | 4.00 | -1 | 16.50 | 0 | | | | | | | | + | 1 | 0.99 | ACP | 0.15 | Non-AntiCP | 0.1882 | Non-ACP | 0.0664 | Non-ACP |
| PAVVLPVPQPEI | Coil | P05814 | CSN2 | Beta-casein | 293 | 53.76 | 12 | 1257.73 | 3.01 | -1 | 9.61 | 0 | | | | + | | | | | 1 | 0.99 | ACP | 0.19 | Non-AntiCP | 0.5786 | ACP | 0.4218 | Non-ACP |
| QELLLNPTHQIYPVTQPLAP VHNPISV | Sheet-Coil-Sheet | P05814 | CSN2 | Beta-casein | 328 | 80.58 | 27 | 3017.63 | 6.05 | -1 | 13.03 | 1490 | | | | | + | | | | 1 | 0.87 | ACP | 0.15 | Non-AntiCP | 0.0782 | Non-ACP | 0.0363 | Non-ACP |
| DEGYGPPPP | Coil | P14866 | HNRNPL | Heterogeneous nuclear ribonucleoprotein L | 154 | 55.70 | 9 | 927.40 | 2.82 | -2 | 17.32 | 1490 | | + | | | | | | | 1 | 0.61 | Non-ACP | 0.37 | Non-AntiCP | 0.4928 | Non-ACP | 0.6799 | ACP |
| SSEESITEYK | Helix | P05814 | CSN2 | Beta-casein | 1276 | 55.19 | 10 | 1171.52 | 3.79 | -2 | 21.39 | 1490 | | + | | | | | | | 1 | 0.99 | ACP | 0.16 | Non-AntiCP | 0.4327 | Non-ACP | 0.0271 | Non-ACP |
| SSSEESITEYK | Helix | P05814 | CSN2 | Beta-casein | 607 | 75.94 | 11 | 1258.55 | 3.79 | -2 | 21.85 | 1490 | + | + | | | | | | | 2 | 0.99 | ACP | 0.18 | Non-AntiCP | 0.4384 | Non-ACP | 0.0304 | Non-ACP |
| EGDFLAEGGGVR | Helix-Coil | P02671 | FGA | Fibrinogen alpha chain | 417 | 69.45 | 12 | 1205.57 | 3.73 | -2 | 22.29 | 0 | | + | | | | | | | 1 | 0.61 | Non-ACP | 0.27 | Non-AntiCP | 0.1798 | Non-ACP | 0.0711 | Non-ACP |
| LSSSEESITEYK | Helix | P05814 | CSN2 | Beta-casein | 1276 | 60.48 | 12 | 1371.64 | 3.80 | -2 | 20.60 | 1490 | | + | + | | | | | | 2 | 0.99 | ACP | 0.22 | Non-AntiCP | 0.3984 | Non-ACP | 0.0293 | Non-ACP |
| LTDLENLHLPLP | Helix-Coil | P05814 | CSN2 | Beta-casein | 293 | 44.08 | 12 | 1373.75 | 4.00 | -2 | 12.63 | 0 | | | | + | | | | | 1 | 0.99 | ACP | 0.19 | Non-AntiCP | 0.5075 | Non-ACP | 0.5333 | Non-ACP |
| FAEEKAVADTRDQ | Helix-Coil | P01833 | PIGR | Polymeric immunoglobulin receptor | 611 | 62.73 | 13 | 1478.70 | 4.00 | -2 | 27.40 | 0 | + | | | | | | | | 1 | 0.93 | ACP | 0.1 | Non-AntiCP | 0.1323 | Non-ACP | 0.0288 | Non-ACP |
| GEGDFLAEGGGVR | Coil-Helix-Coil | P02671 | FGA | Fibrinogen alpha chain | 417 | 89.82 | 13 | 1262.59 | 3.74 | -2 | 23.44 | 0 | | + | | | | | | | 1 | 0.98 | ACP | 0.23 | Non-AntiCP | 0.1951 | Non-ACP | 0.2045 | Non-ACP |
| SLSSSEESITEYK | Helix | P05814 | CSN2 | Beta-casein | 1276 | 58.12 | 13 | 1458.67 | 3.79 | -2 | 21.06 | 1490 | | + | | | | | | | 1 | 0.99 | ACP | 0.21 | Non-AntiCP | 0.4402 | Non-ACP | 0.0549 | Non-ACP |
| SGEGDFLAEGGGVR | Coil-Helix-Coil | P02671 | FGA | Fibrinogen alpha chain | 243 | 103.90 | 14 | 1349.62 | 3.73 | -2 | 23.90 | 0 | + | + | + | | | | | | 3 | 0.99 | ACP | 0.14 | Non-AntiCP | 0.1650 | Non-ACP | 0.0248 | Non-ACP |
| LFAEEKAVADTRDQA | Helix | P01833 | PIGR | Polymeric immunoglobulin receptor | 611 | 58.22 | 15 | 1662.82 | 4.00 | -2 | 26.65 | 0 | + | | | + | | | | | 2 | 0.97 | ACP | 0.07 | Non-AntiCP | 0.1193 | Non-ACP | 0.0274 | Non-ACP |
| RLQNPSESSEPIPLE | Coil | P47710 | CSN1S1 | Alpha-S1-casein | 62 | 60.06 | 15 | 1694.84 | 3.78 | -2 | 20.40 | 0 | | + | | | | | | | 1 | 0.99 | ACP | 0.07 | Non-AntiCP | 0.2424 | Non-ACP | 0.0372 | Non-ACP |
| DSGSSEEQGGSSRALV | Helix-Coil | P01833 | PIGR | Polymeric immunoglobulin receptor | 956 | 54.91 | 16 | 1564.69 | 3.73 | -2 | 25.92 | 0 | | + | | | | | | | 1 | 0.86 | Non-ACP | 0.13 | Non-AntiCP | 0.3597 | Non-ACP | 0.0289 | Non-ACP |
| SGVENALTKSELLVEQ | Helix | Q99541 | PLIN2 | Perilipin-2 | 58 | 58.12 | 16 | 1715.89 | 3.79 | -2 | 21.36 | 0 | | + | | | | | | | 1 | 0.97 | ACP | 0.02 | Non-AntiCP | 0.0854 | Non-ACP | 0.0311 | Non-ACP |
| SVDSGSSEEQGGSSRA | Coil | P01833 | PIGR | Polymeric immunoglobulin receptor | 956 | 85.81 | 16 | 1538.64 | 3.73 | -2 | 27.63 | 0 | | + | | | | | | | 1 | 0.99 | ACP | 0.2 | Non-AntiCP | 0.4645 | Non-ACP | 0.0385 | Non-ACP |
| SRASVDSGSSEEQGGSS | Coil-Helix-Coil | P01833 | PIGR | Polymeric immunoglobulin receptor | 956 | 53.69 | 17 | 1625.67 | 3.73 | -2 | 28.09 | 0 | | + | | | | | | | 1 | 0.99 | ACP | 0.19 | Non-AntiCP | 0.5173 | ACP | 0.0434 | Non-ACP |
| SVDSGSSEEQGGSSRAL | Coil | P01833 | PIGR | Polymeric immunoglobulin receptor | 956 | 55.48 | 17 | 1651.73 | 3.73 | -2 | 26.38 | 0 | | + | | | | | | | 1 | 0.97 | ACP | 0.16 | Non-AntiCP | 0.3947 | Non-ACP | 0.0291 | Non-ACP |

| | | | | | | | | | | | | | | | | | | | | | | | | | | |
|---|---|---|---|---|---|---|---|---|---|---|---|---|---|---|---|---|---|---|---|---|---|---|---|---|---|---|
| FAEEKAVADTRDQADGSR | Helix-Coil | P01833 | PIGR | Polymeric immunoglobulin receptor | 93 | 62.20 | 18 | 1964.92 | 4.15 | -2 | 34.96 | 0 | | | | + | | 1 | 0.98 | ACP | 0.08 | Non-AntiCP | 0.0697 | Non-ACP | 0.0589 | Non-ACP |
| LFAEEKAVADTRDQADGSR | Helix-Coil | P01833 | PIGR | Polymeric immunoglobulin receptor | 956 | 66.70 | 19 | 2078.00 | 4.16 | -2 | 33.71 | 0 | + | + | + | | | 3 | 0.99 | ACP | 0.06 | Non-AntiCP | 0.0566 | Non-ACP | 0.0412 | Non-ACP |
| AIQDPRLFAEEKAVADTRDQ | Coil-Helix-Coil | P01833 | PIGR | Polymeric immunoglobulin receptor | 93 | 40.52 | 20 | 2272.14 | 4.16 | -2 | 31.89 | 0 | | | + | + | | 2 | 0.86 | Non-ACP | 0.03 | Non-AntiCP | 0.0578 | Non-ACP | 0.0354 | Non-ACP |
| ADGSRASVDSGSSEEQGGSSR | Coil | P01833 | PIGR | Polymeric immunoglobulin receptor | 956 | 60.40 | 21 | 2024.86 | 4.00 | -2 | 35.19 | 0 | | + | | | | 1 | 0.61 | Non-ACP | 0.24 | Non-AntiCP | 0.4106 | Non-ACP | 0.0358 | Non-ACP |
| DSGSSEEQGGSSRALVSTLVPLG | Helix-Coil | P01833 | PIGR | Polymeric immunoglobulin receptor | 93 | 59.03 | 23 | 2232.08 | 3.73 | -2 | 24.96 | 0 | + | | | + | | 2 | 0.86 | Non-ACP | 0.07 | Non-AntiCP | 0.1669 | Non-ACP | 0.0344 | Non-ACP |
| ASVDSGSSEEQGGSSRALVSTLVPLG | Coil-Helix-Coil | P01833 | PIGR | Polymeric immunoglobulin receptor | 93 | 32.47 | 26 | 2489.22 | 3.74 | -2 | 25.46 | 0 | | | | + | | 1 | 0.86 | Non-ACP | 0.03 | Non-AntiCP | 0.1162 | Non-ACP | 0.0706 | Non-ACP |
| DAAPDEKVLDSGF | Helix-Coil | P01833 | PIGR | Polymeric immunoglobulin receptor | 956 | 57.27 | 13 | 1362.63 | 3.48 | -3 | 24.58 | 0 | | + | | | | 1 | 0.80 | Non-ACP | 0.25 | Non-AntiCP | 0.1002 | Non-ACP | 0.0356 | Non-ACP |
| DGREQEAEQMPEYR | Helix-Coil | Q13410 | BTN1A1 | Butyrophilin subfamily 1 member A1 | 186 | 60.41 | 14 | 1736.74 | 3.87 | -3 | 31.63 | 1490 | | + | | | | 1 | 0.86 | Non-ACP | 0.23 | Non-AntiCP | 0.3099 | Non-ACP | 0.0433 | Non-ACP |
| ESLSSSEESITEYK | Helix | P05814 | CSN2 | Beta-casein | 607 | 59.51 | 14 | 1587.71 | 3.61 | -3 | 24.69 | 1490 | + | + | | | | 2 | 0.61 | Non-ACP | 0.26 | Non-AntiCP | 0.4352 | Non-ACP | 0.0367 | Non-ACP |
| RETIESLSSSEESI | Helix | P05814 | CSN2 | Beta-casein | 1276 | 54.61 | 14 | 1566.74 | 3.61 | -3 | 23.29 | 0 | | + | | | | 1 | 0.99 | ACP | 0.1 | Non-AntiCP | 0.4268 | Non-ACP | 0.0637 | Non-ACP |
| AEEKAVADTRDQADG | Helix | P01833 | PIGR | Polymeric immunoglobulin receptor | 956 | 56.18 | 15 | 1574.71 | 3.81 | -3 | 34.40 | 0 | | + | | | | 1 | 0.61 | Non-ACP | 0.1 | Non-AntiCP | 0.1380 | Non-ACP | 0.0475 | Non-ACP |
| DGREQEAEQMPEYRG | Helix-Coil | Q13410 | BTN1A1 | Butyrophilin subfamily 1 member A1 | 186 | 57.93 | 15 | 1793.76 | 3.87 | -3 | 32.78 | 1490 | | + | | | | 1 | 0.86 | Non-ACP | 0.18 | Non-AntiCP | 0.2364 | Non-ACP | 0.0434 | Non-ACP |
| DSGEGDFLAEGGGVR | Coil-Helix-Coil | P02671 | FGA | Fibrinogen alpha chain | 417 | 105.48 | 15 | 1464.65 | 3.52 | -3 | 27.54 | 0 | + | + | + | | | 3 | 0.61 | Non-ACP | 0.11 | Non-AntiCP | 0.1222 | Non-ACP | 0.0346 | Non-ACP |
| IESLSSSEESITEYK | Helix | P05814 | CSN2 | Beta-casein | 1276 | 76.40 | 15 | 1700.80 | 3.62 | -3 | 23.57 | 1490 | + | + | + | | | 3 | 0.99 | ACP | 0.2 | Non-AntiCP | 0.4110 | Non-ACP | 0.0355 | Non-ACP |
| ADSGEGDFLAEGGGVR | Helix-Coil | P02671 | FGA | Fibrinogen alpha chain | 417 | 99.99 | 16 | 1535.68 | 3.54 | -3 | 28.04 | 0 | + | + | | | + | 3 | 0.61 | Non-ACP | 0.16 | Non-AntiCP | 0.1013 | Non-ACP | 0.0286 | Non-ACP |
| AEEKAVADTRDQADGS | Helix | P01833 | PIGR | Polymeric immunoglobulin receptor | 611 | 66.13 | 16 | 1661.75 | 3.81 | -3 | 34.86 | 0 | + | + | | | | 2 | 0.61 | Non-ACP | 0.08 | Non-AntiCP | 0.0990 | Non-ACP | 0.0405 | Non-ACP |
| FAEEKAVADTRDQADG | Helix | P01833 | PIGR | Polymeric immunoglobulin receptor | 956 | 58.74 | 16 | 1721.78 | 3.80 | -3 | 32.69 | 0 | | + | | | | 1 | 0.98 | ACP | 0.12 | Non-AntiCP | 0.1123 | Non-ACP | 0.0341 | Non-ACP |
| TIESLSSSEESITEYK | Helix | P05814 | CSN2 | Beta-casein | 1276 | 63.79 | 16 | 1801.84 | 3.61 | -3 | 23.82 | 1490 | | + | | | | 1 | 0.99 | ACP | 0.16 | Non-AntiCP | 0.3726 | Non-ACP | 0.0363 | Non-ACP |
| FAEEKAVADTRDQADGS | Helix | P01833 | PIGR | Polymeric immunoglobulin receptor | 956 | 63.11 | 17 | 1808.81 | 3.80 | -3 | 33.15 | 0 | | + | + | | | 2 | 0.98 | ACP | 0.11 | Non-AntiCP | 0.0819 | Non-ACP | 0.0427 | Non-ACP |
| SAWLDSGVTGSGLEGDH | Helix-Coil | P12272 | PTHLH | Parathyroid hormone-related protein | 70 | 70.36 | 17 | 1686.75 | 3.68 | -3 | 22.82 | 5500 | | + | | | | 1 | 0.99 | ACP | 0.32 | Non-AntiCP | 0.1388 | Non-ACP | 0.1174 | Non-ACP |
| RETIESLSSSEESITEYK | Helix | P05814 | CSN2 | Beta-casein | 1276 | 77.19 | 18 | 2086.99 | 3.90 | -3 | 29.26 | 1490 | | + | + | | | 2 | 0.99 | ACP | 0.15 | Non-AntiCP | 0.3007 | Non-ACP | 0.0396 | Non-ACP |
| ADGSRASVDSGSSEEQGGSS | Coil | P01833 | PIGR | Polymeric immunoglobulin receptor | 956 | 70.74 | 20 | 1868.76 | 3.54 | -3 | 33.38 | 0 | | + | | | | 1 | 0.61 | Non-ACP | 0.17 | Non-AntiCP | 0.4438 | Non-ACP | 0.0445 | Non-ACP |
| DQADGSRASVDSGSSEEQGGSSR | Coil-Helix-Coil | P01833 | PIGR | Polymeric immunoglobulin receptor | 956 | 80.94 | 23 | 2267.95 | 3.80 | -3 | 39.60 | 0 | | + | | | | 1 | 0.61 | Non-ACP | 0.18 | Non-AntiCP | 0.3058 | Non-ACP | 0.0440 | Non-ACP |
| DQADGSRASVDSGSSEEQGGSSRA | Coil-Helix-Coil | P01833 | PIGR | Polymeric immunoglobulin receptor | 956 | 65.97 | 24 | 2338.98 | 3.80 | -3 | 40.10 | 0 | | + | | | | 1 | 0.61 | Non-ACP | 0.14 | Non-AntiCP | 0.2701 | Non-ACP | 0.0450 | Non-ACP |
| VADTRDQADGSRASVDSGSSEEQGGSSRA | Coil | P01833 | PIGR | Polymeric immunoglobulin receptor | 611 | 74.95 | 29 | 2881.26 | 3.97 | -3 | 45.84 | 0 | + | + | | | | 2 | 0.99 | ACP | 0.06 | Non-AntiCP | 0.1182 | Non-ACP | 0.0395 | Non-ACP |
| RPDIQYPDATDEDIT | Coil | P10451 | SPP1 | Osteopontin | 159 | 74.59 | 15 | 1747.79 | 3.35 | -4 | 27.00 | 1490 | | + | | | | 1 | 0.99 | ACP | 0.08 | Non-AntiCP | 0.2386 | Non-ACP | 0.0471 | Non-ACP |

| | | | | | | | | | | | | | | | | | | | | | | | | | | | | | | | |
|---|---|---|---|---|---|---|---|---|---|---|---|---|---|---|---|---|---|---|---|---|---|---|---|---|---|---|---|---|---|---|---|
| PPPPEPVPMYSDEDLA | Coil | O60318 | MCM3AP | Germinal-center associated nuclear protein | 48 | 48.05 | 16 | 1752.79 | 2.64 | -4 | 21.15 | 1490 | | | | + | | | | | | | 1 | 0.97 | ACP | 0.19 | Non-AntiCP | 0.2262 | Non-ACP | 0.1044 | Non-ACP |
| RETIESLSSSEESITE | Helix | P05814 | CSN2 | Beta-casein | 1276 | 67.27 | 16 | 1795.83 | 3.48 | -4 | 27.17 | 0 | + | + | | | | | | | | | 2 | 0.99 | ACP | 0.15 | Non-AntiCP | 0.3523 | Non-ACP | 0.0617 | Non-ACP |
| ETIESLSSSEESITEYK | Helix | P05814 | CSN2 | Beta-casein | 293 | 74.50 | 17 | 1930.89 | 3.49 | -4 | 27.45 | 1490 | + | + | + | + | + | + | | | | + | 7 | 0.61 | Non-ACP | 0.17 | Non-AntiCP | 0.3562 | Non-ACP | 0.0372 | Non-ACP |
| RETIESLSSSEESITEY | Helix | P05814 | CSN2 | Beta-casein | 1276 | 82.63 | 17 | 1958.89 | 3.48 | -4 | 26.46 | 1490 | + | + | + | | | | | | | | 3 | 0.99 | ACP | 0.13 | Non-AntiCP | 0.3355 | Non-ACP | 0.0561 | Non-ACP |
| RPDIQYPDATDEDITSH | Helix-Coil | P10451 | SPP1 | Osteopontin | 159 | 76.52 | 17 | 1971.88 | 3.76 | -4 | 29.79 | 1490 | | + | | | | | | | | | 1 | 0.99 | ACP | 0.09 | Non-AntiCP | 0.1628 | Non-ACP | 0.0499 | Non-ACP |
| ETIESLSSSEESITEYKQ | Helix | P05814 | CSN2 | Beta-casein | 1276 | 63.19 | 18 | 2058.94 | 3.49 | -4 | 28.22 | 1490 | | + | | | | | | | | | 1 | 0.61 | Non-ACP | 0.1 | Non-AntiCP | 0.3281 | Non-ACP | 0.0425 | Non-ACP |
| VKHEDQQQGEDEHQDKIYP | Helix-Coil-Helix | P05814 | CSN2 | Beta-casein | 467 | 62.91 | 19 | 2322.05 | 4.29 | -4 | 42.05 | 1490 | | | + | | | | | | | | 1 | 0.97 | ACP | 0.12 | Non-AntiCP | 0.1370 | Non-ACP | 0.0435 | Non-ACP |
| DQADGSRASVDSGSSEEQGGSS | Coil-Helix-Coil | P01833 | PIGR | Polymeric immunoglobulin receptor | 611 | 79.76 | 22 | 2111.84 | 3.39 | -4 | 37.79 | 0 | + | + | | | | | | | | | 2 | 0.61 | Non-ACP | 0.17 | Non-AntiCP | 0.3397 | Non-ACP | 0.0450 | Non-ACP |
| AVADTRDQADGSRASVDSGSSEE | Helix-Coil | P01833 | PIGR | Polymeric immunoglobulin receptor | 956 | 71.47 | 23 | 2309.00 | 3.67 | -4 | 40.04 | 0 | + | + | + | | | | | | | | 3 | 0.61 | Non-ACP | 0.05 | Non-AntiCP | 0.0769 | Non-ACP | 0.0540 | Non-ACP |
| AVADTRDQADGSRASVDSGSSEEQ | Helix-Coil | P01833 | PIGR | Polymeric immunoglobulin receptor | 956 | 74.88 | 24 | 2437.06 | 3.67 | -4 | 40.81 | 0 | + | + | + | | | | | | | | 3 | 0.86 | Non-ACP | 0.04 | Non-AntiCP | 0.0598 | Non-ACP | 0.0542 | Non-ACP |
| VADTRDQADGSRASVDSGSSEEQG | Coil-Helix-Coil | P01833 | PIGR | Polymeric immunoglobulin receptor | 956 | 64.42 | 24 | 2423.04 | 3.67 | -4 | 41.46 | 0 | | + | | | | | | | | | 1 | 0.99 | ACP | 0.05 | Non-AntiCP | 0.0667 | Non-ACP | 0.0486 | Non-ACP |
| AVADTRDQADGSRASVDSGSSEEQG | Helix-Coil | P01833 | PIGR | Polymeric immunoglobulin receptor | 956 | 60.32 | 25 | 2494.08 | 3.67 | -4 | 41.96 | 0 | + | + | | | | | | | | | 2 | 0.61 | Non-ACP | 0.07 | Non-AntiCP | 0.0613 | Non-ACP | 0.0527 | Non-ACP |
| ADTRDQADGSRASVDSGSSEEQGGSS | Helix-Coil-Helix | P01833 | PIGR | Polymeric immunoglobulin receptor | 956 | 63.51 | 26 | 2555.06 | 3.67 | -4 | 43.99 | 0 | | + | | | | | | | | | 1 | 0.61 | Non-ACP | 0.08 | Non-AntiCP | 0.1933 | Non-ACP | 0.0373 | Non-ACP |
| VADTRDQADGSRASVDSGSSEEQGGSS | Coil-Helix-Coil | P01833 | PIGR | Polymeric immunoglobulin receptor | 956 | 81.48 | 27 | 2654.12 | 3.67 | -4 | 43.53 | 0 | + | + | | | | | | | | | 2 | 0.99 | ACP | 0.05 | Non-AntiCP | 0.1313 | Non-ACP | 0.0387 | Non-ACP |
| AVADTRDQADGSRASVDSGSSEEQGGSS | Coil | P01833 | PIGR | Polymeric immunoglobulin receptor | 956 | 93.07 | 28 | 2725.16 | 3.67 | -4 | 44.03 | 0 | + | + | + | | | | | | | | 3 | 0.61 | Non-ACP | 0.07 | Non-AntiCP | 0.1145 | Non-ACP | 0.0426 | Non-ACP |
| ETIESLSSSEESITE | Helix | P05814 | CSN2 | Beta-casein | 607 | 60.39 | 15 | 1639.73 | 2.70 | -5 | 25.36 | 0 | + | | | | | | | | | | 1 | 0.61 | Non-ACP | 0.16 | Non-AntiCP | 0.4012 | Non-ACP | 0.1003 | Non-ACP |

NA: not applicable
ACP: anti-cancer peptide
NP: non-prediction
* predicted by PepDraw tool (http://www.tulane.edu/~biochem/WW/PepDraw/)

**Table S3**. Physicochemical, structural and machine learning properties of the selected peptides.

| **Peptide name** | **Peptide sequence** | **Length** | **Mass (Da)** | **pI** | **Net charge** | **Structure*** | **Anticancer peptide prediction (predictive value) (Y= anticancer peptide; N= non-anticancer peptide)** | | | |
|---|---|---|---|---|---|---|---|---|---|---|
| | | | | | | | **ACPred-FL** | **AntiCP 2.0** | **MLACP** | **mACPpred** |
| HMP-S1 | ETIESLSSSEESITEYK | 17 | 1932.26 | 3.99 | -4 | Helix | N(60.63%) | N(0.15) | N(0.3562) | N(0.0372) |
| HMP-S2 | ADSGEGDFLAEGGGVR | 16 | 1693.00 | 3.92 | -3 | Coil-Helix-Coil | N(60.63%) | N(0.16) | N(0.1013) | N(0.0286) |
| HMP-S3 | PPPPPPPPP | 9 | 989.30 | 5.88 | 0 | Coil | Y(81.52%) | Y(0.67) | Y(0.6377) | Y(0.9287) |
| HMP-S4 | APGPP | 5 | 534.68 | 5.88 | 0 | Coil | NP | Y(0.60) | Y(0.5979) | Y(0.9276) |
| HMP-S5 | VSLISAEVPLGR | 12 | 1396.84 | 6.36 | 0 | Coil-Helix | Y(99.22%) | N(0.14) | N(0.3000) | N(0.0435) |
| BMP-S6 (positive control) | FKCRRWQWRMKKLGAPSITCVR (*48*) | 22 | 2751.69 | 11.58 | +7 | Coil-Helix | N(60.63%) | Y(1.0) | Y(0.9444) | Y(0.9848) |
| HMP-S7 | SFIPRAKSTWLNNIKLL | 17 | 2114.84 | 11.17 | +3 | Helix | Y(94.40%) | Y(0.57) | N(0.4010) | Y(0.8932) |
| HMP-S8 | GRATLVQDGIAKGRVA | 16 | 1683.19 | 10.84 | +2 | Helix-Coil | Y(83.42%) | N(0.25) | N(0.0824) | N(0.2289) |
| HMP-S9 | LPIPQQVVPYPQRAVPVQ | 18 | 2157.84 | 9.10 | +1 | Coil-Sheet | Y(99.22%) | N(0.33) | N(0.4242) | N(0.0783) |

NP: not predictable; HMP: human milk peptide; BMP: bovine milk peptide
*: predicted by PEP-FOLD3 software
